# Deferoxamine as adjunct therapeutics for tuberculosis

**DOI:** 10.1101/2023.06.02.543389

**Authors:** Sandeep R. Kaushik, Nidhi Yadav, Ashish Gupta, Sukanya Sahu, Nikhil Bhalla, Poonam Dagar, Amit Kumar Mohapatra, Adyasha Sarangi, Taruna Sharma, Bichitra Biswal, Amit Kumar Pandey, Ranjan Kumar Nanda

## Abstract

Dysregulated iron metabolism is reported in tuberculosis patients; therefore, it represents an opportunity for developing host-directed therapeutics. This study monitored the antimycobacterial properties of an iron chelator, i.e., Deferoxamine (DFO/D), and its impact on the transcript and metabolite levels of *Mycobacterium tuberculosis* (Mtb) *in vitro*. For *in vivo* validation, a group of mice received ferric carboxymaltose to create an iron overload condition, and controls were aerosol-infected with Mtb H37Rv. Mtb-infected mice received isoniazid (INH/H) and rifampicin (RIF/R) in combination with or without DFO before tissue-specific CFU assay, liver metabolite screening and iron quantification using mass spectrometry. DFO showed antimycobacterial properties comparable to INH *in vitro*. DFO treatment deregulated (log2DFO/control>±1.0) Mtb transcript (n=137) levels, the majority of which encode for iron-containing proteins and proteins involved in stress response. DFO treatment up-regulated Rv3622c (PE32), Rv2353c (PPE39) and Rv3022A (PE29) genes and conditional knocking down of ABC transporter like *irtA* by anhydrotetracycline (Atc) inducible CRISPR interference (CRISPRi) approach compromised Mtb growth showing their potential involvement in iron metabolism. Global Mtb metabolite analysis using GC-MS identified a set of 5 deregulated metabolites indicating a perturbed pentose phosphate pathway and inositol phosphate metabolism in the host upon DFO treatment. Iron-overloaded mice exhibited significantly higher tissue mycobacterial burden at two weeks post-infection, and the efficacy of INH and RIF were compromised, corroborating with previous reports. Iron chelation by DFO or combined with/adjunct to RIF and INH significantly reduced the lung/tissue mycobacterial burden at four weeks post-treatment, specifically in the first (∼0.5 log) and second weeks (∼0.5 log) of treatment. The intracellular pro-inflammatory cytokine levels in the lung CD4+ T cells of INH and RIF-treated groups with or without DFO were similar, suggesting DFO has a direct role in Mtb survival and metabolism rather than improved infection, and the efficacy of INH and RIF were compromised, corroborating with previous reports. DFO adjunct to RIF and INH treatment significantly altered liver arginine biosynthesis, which directly neutralizes ammonia and is immune-supportive. Conventionally, DFO is used for treating acute iron toxicity that is common in thalassemic patients, and this study demonstrates DFO has potential as adjunct therapeutics for tuberculosis.

**Graphical abstract:** 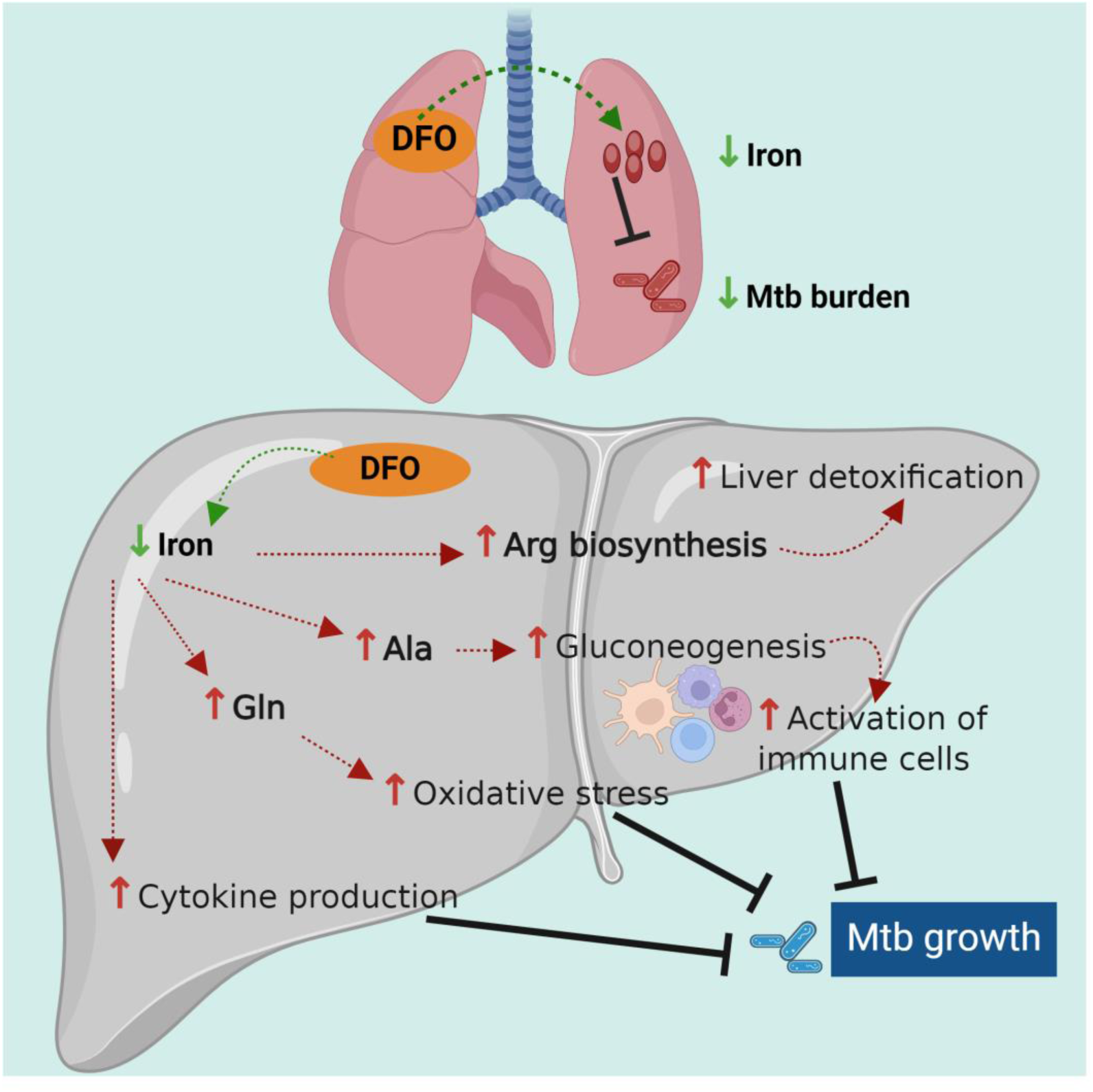

## Introduction

*Mycobacterium tuberculosis* (Mtb), the causative agent of Tuberculosis (TB), primarily infects the lower respiratory tract. After reaching the lungs, through the lymphatic and immune cell system, it disseminates to other body parts, causing Miliary and extra-pulmonary TB in different organs like the urinogenital system, brain, spinal cord, and liver.^1^ The duration of treatment modalities for TB patients infected with drug-sensitive Mtb strains is 6–9 months and may prolong up to 24 months or more if drug-resistant Mtb strains cause the infection.^2^ Shortening TB treatment duration by adopting additional host-directed therapeutics might help in increasing adherence and may also reduce the relapse rate, minimize Mtb drug resistance development, and may help in achieving the eradication of TB.^3^

Iron is an essential micronutrient for the host and Mtb’s survival. In the host, iron is absorbed from the food materials through the intestine and is involved in erythropoiesis, respiration and other essential functions. Meanwhile, Mtb, with the help of siderophores like mycobactin and carboxymycobactin, chelates the host Fe^3+^ ion and intracellularly stores it in Fe^2+^ form with the help of iron storage proteins.^4,5^ Iron supplementation in TB patients is reported to worsen clinical outcomes. In contrast, iron chelation in *in vitro* Mtb cultures prevents its growth and significantly reduces Mtb load in *in vivo* settings.^6,7,8,9,10^ Infecting mice or macrophages with Mtb strain deficient in synthesis or uptake of siderophores leads to an inefficient establishment of active infection.^4^ Mtb experiences iron deprivation in the endosome after being engulfed by interstitial macrophages. This compartment of interstitial macrophages is not permissive for Mtb growth. However, in alveolar macrophages, Mtb seldom experiences iron deprivation, and therefore, iron is readily available for Mtb inside its endosome.^11^ Iron deprivation in Mtb compromises functions of enzymes that need iron as cofactors as those involved in the TCA cycle and oxidative phosphorylation; therefore, its deprivation is known to cause metabolic reprogramming in the host.^12^

The actinomycetes *Streptomyces pilosus* produces deferoxamine (DFO), that acts as a siderophore, having strong affinity towards iron (Fe^3+^), and forms a stable ferrioxamine complex. DFO is hydrophilic and is used for the treatment of iron-toxicity which is associated with various conditions including hemochromatosis. Excess iron *in vitro* increases the viability and proliferation of Mtb, whereas chelation using DFO affects both these activities, whereas iron chelation by silybin only impacts Mtb proliferation.^13^ Chelating host iron by DFO may also deprive Mtb from iron, which is essential for its survival and establishment of the active TB disease.

In this study, we aimed to elucidate the anti-Mtb potential of DFO by monitoring its influence on Mtb survival, levels of transcripts involved in iron metabolism, impact on tissue metabolism and whether at early treatment time points it accelerates Mtb clearance adjunct to the existing anti-TB drugs in C57BL/6 mice. Our findings indicate that iron deprivation caused by DFO can augment earlier Mtb clearance from the host.

## Results

### Deferoxamine affects mycobacterial growth *in vitro*

Following the existing literature, the DFO dose was fixed and used in all our experiments.^14,15^ The growth curve of Mtb showed a time-dependent increase in controls, whereas replicates treated with DFO at 0.1 mg/mL or INH at 0.1 μg/mL alone or combined compromised Mtb growth (Figure 1A and B). The growth kinetics of Mtb in the presence of DFO in 7H9 media were comparable to INH-treated groups (Figure 1B). Furthermore, the combined treatment of DFO and INH also compromised Mtb growth. On day five, post-inoculation, a significant reduction in Mtb survival was observed in the presence of DFO, INH as well as their combination (Figure 1C). However, as expected, DFO alone or in combination with INH significantly reduced intracellular iron, zinc and copper levels in Mtb (Figure 1D, Supplementary Figure S1A and B). From this we concluded that DFO may have a bactericidal effect, as Mtb relies on these divalent ions for various processes like two-metal ion catalysis that are essential for its survival and growth.

**Figure 1:**
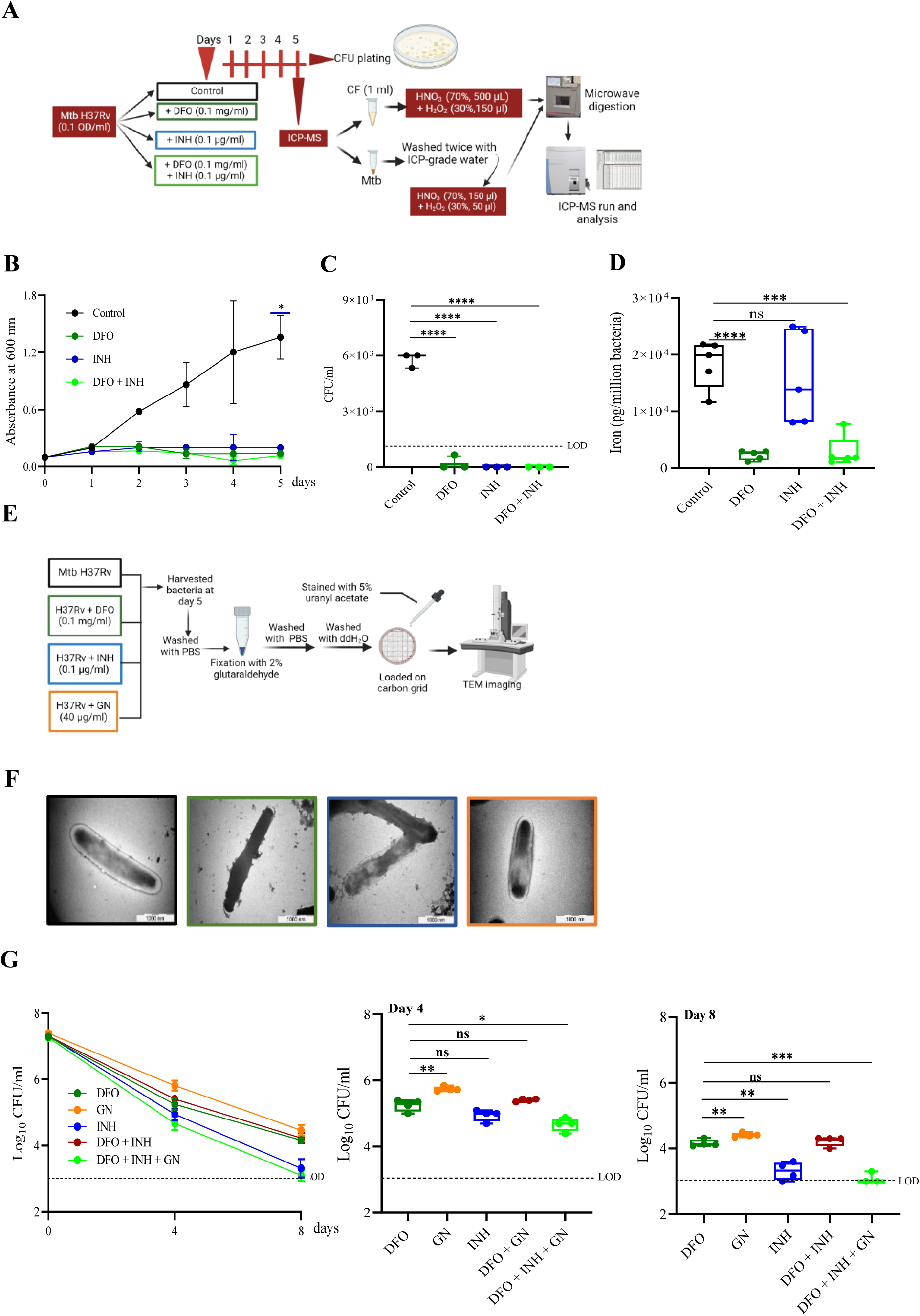
Deferoxamine (DFO) chelates iron and shows mycobactericidal effect in the *in-vitro* condition. **A.** Schematic presentation of the method used for *in vitro* Mtb culture with DFO and isoniazid (INH) for growth kinetics and ICP-MS. **B.** Growth kinetics of *Mycobacterium tuberculosis* H37Rv strain in the absence and presence of different drugs (DFO, INH, DFO+INH) and control. **C.** Survived Mycobacteria cultures on day five post-drug-treated conditions and control as observed in CFU assay, showing the significant bactericidal effect of the drugs alone or in combination. **D.** Intracellular elemental levels in drug-treated and controlled mycobacterial culture at 5th-day post-treatment quantified using ICP-MS. **E.** Schematic presentation of the methodology used for TEM sample preparation. **F.** Transmission electron micrographs of untreated, DFO, INH, and Gallium nitrate (GN) treated Mtb, respectively, harvested at the 5th day post-drug treatment. **G.** Kill curve of Mtb with DFO, GN, INH and combination at 10x MIC of each drug. **H.** CFU plot for day four post-inoculation. CFU plot for day eight post-inoculation. CF: culture filtrate, CFU: colony forming units, and dashed line in the plots represent the limit of detection (LOD). Each time point had five biological replicates per group. Results are determined using unpaired, non-parametric two-tailed t-tests. ns: not significant. ns: not significant; *: p<0.05; **: p<0.01; ***: p<0.001; ****: p<0.005 at 95% confidence interval.

Because Gallium nitrate is reported to have antimycobacterial properties, we used it as an additional control at 40 μg/mL concentration in subsequent experiments.^21^ Electron micrographs of Mtb treated with DFO and INH-RIF, harvested on day five post-treatment, showed changes in the cell wall compared to the control and Gallium nitrate (GN) treated Mtb (Figure 1E and F, Supplementary Figure S2). The kill kinetics of Mtb in the presence of DFO were comparable to INH at day four and in combination (DFO+GN+INH) had significantly lower CFU count at day eight post-incubation, whereas GN in combination with DFO showed comparable CFU at both day 4 and 8 (Figure 1G). GN exhibited a less pronounced anti-Mtb potential than DFO and INH, so it was excluded from further experiments. These results revealed that DFO restricts iron availability, reduces CFU count, has anti-Mtb potential, and causes morphological changes in Mtb.

### DFO limits iron, leading to deregulated iron metabolism in Mtb

To understand the direct impact of DFO on the Mtb transcriptome machinery, RNA-seq analysis of Mtb treated with DFO and controls was performed. Quality control of RNA-seq data showed ∼18.2±1.6 million high-quality reads based on a Phred-score ≥ 30 with negligible adapter contamination. Upon alignment of the reads with the reference Mtb genome, we observed alignment rates of 74.3±4.1 % and 97.1±0.8 % of DFO-treated and control Mtb cultures, respectively. Out of a set of 3,931 identified Mtb genes, 3,826 showed a counts/million (CPM) value ≥ 1 in all replicates of each condition and 137 genes were deregulated (log_2_DFO/control>±1.0, adjusted p-value ≤ 0.05) (Figure 2A, B, and C). The top 10 down regulated genes included Rv1593c, Rv1596, Rv0263c, Rv1595, Rv0757, Rv1594, Rv0264c, Rv0262c, Rv1130, and Rv1131. The top 10 up regulated genes included Rv0758, Rv2353c, Rv0094c, Rv2812, Rv1587c, Rv2813, Rv1553, Rv1290A, Rv1917c, and Rv2370c (Figure 2C). Principal component analysis (PCA) of all the identified transcripts and unsupervised hierarchical clustering of the deregulated gene sets showed two distinct clusters of the DFO-treated and control samples (Figure 2D, Supplementary Figure S3A). In this study, Mtb treated with DFO showed up-regulation of Rv1094, Rv1594 and the down-regulation of Rv0327, Rv1553, Rv2276, Rv3251c, Rv3260c, and Rv3406 supporting earlier reports (Figure 2E).^16,17^

**Figure 2:**
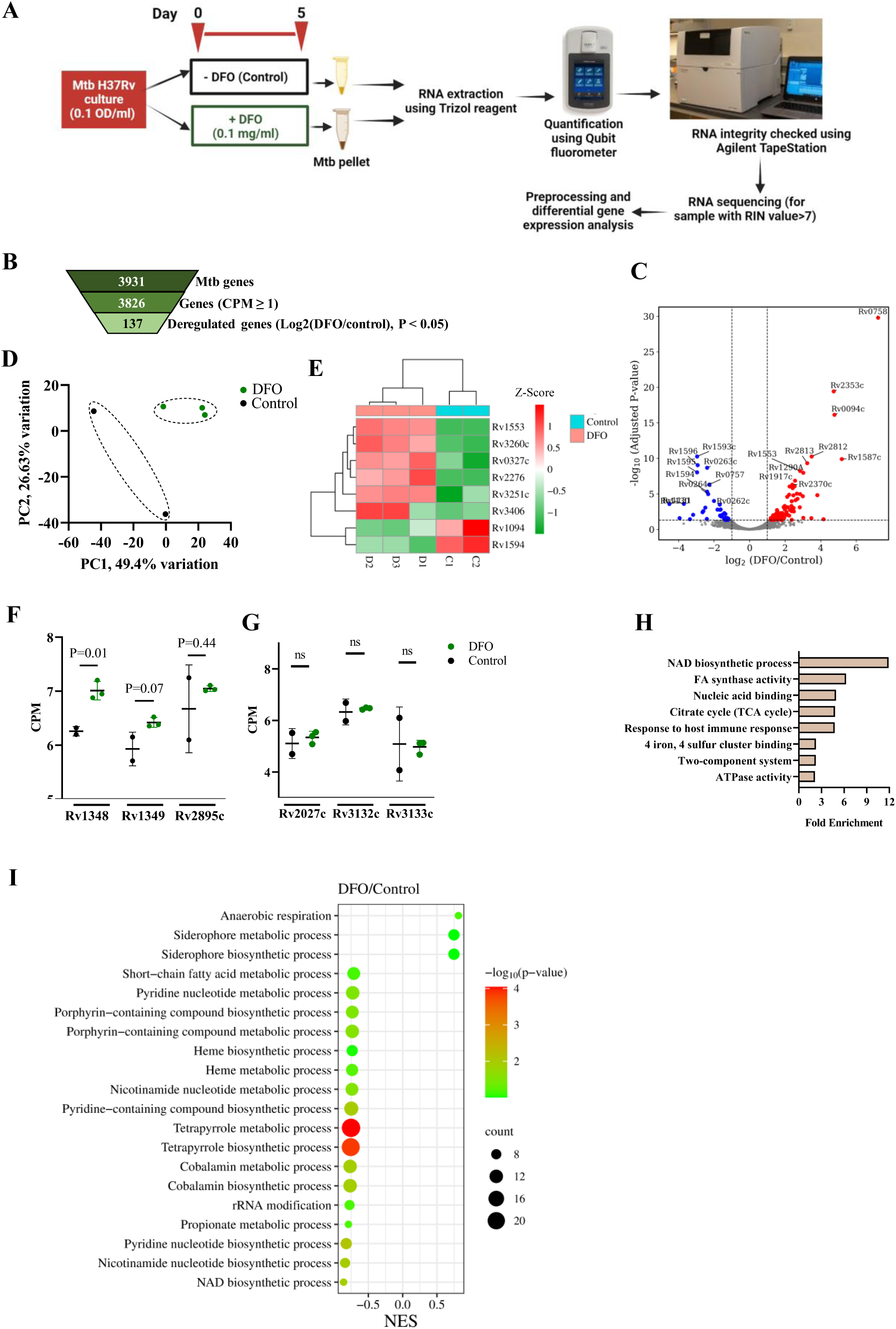
Transcriptome analysis of DFO-treated Mtb showed deregulated iron metabolism and several other pathways. **A.** Schematic presentation of the method used for RNA extraction from DFO-treated Mtb H37Rv and control for RNAseq analysis. **B.** Identified transcripts with CPM ≥ 1, from the RNA-seq analysis and the deregulated (log_2_DFO/control ≥ ± 1.0, p < 0.05) ones. **C.** Volcano plot showing the deregulated genes in DFO-treated Mtb compared to control. **D.** Principal component analysis (PCA) of the RNA-seq data (transcripts: 3826) of DFO treated Mtb and control **E.** Heatmap showing transcript levels of iron deprivation marker genes of DFO treated Mtb and control. **F.** Transcript levels of known iron transporters (Rv1348, Rv1349) and Mycobactin protein Utilization ViuB (Rv2895c) in DFO-treated Mtb and control. **G.** Transcript levels of dormancy-inducing genes *dosT* (Rv2027c), *dosS* (Rv3132) and *dosR* (Rv3133) in DFO-treated Mtb and controls. **H.** Gene Ontology analysis of the deregulated transcripts using the DAVID tool in DFO-treated Mtb. **I.** Deregulated biological processes, as identified from the pre-ranked FGSEA (iDEP.96 tool) in DFO-treated Mtb compared to control.

Iron availability impacts the levels of transporters like Rv1348 (*irtA*), Rv1349 (*irtB*) and mycobactin utilization protein Rv2895c (*viuB*). Since *irtAB* functions as an ABC transporter complex involved in iron scavenging, it’s essential role in iron transport was verified by creating a conditional knockdown strain utilizing an anhydrotetracycline (Atc) inducible CRISPR interference (CRISPRi) approach. A scrambled sgRNA (single guide RNA) was used as a control to assess the gene-specific effect. The suppression of both the genes (*irtA* and *irtB*) in the presence of Atc in the CRISPR knockdown strain of *IrtA* was observed (Supplementary Figure S4A). Interestingly, the Atc-induced *irtA-*KD Mtb did not grow in 7H9 media, confirming its essentiality for its growth and multiplication (Supplementary Figure S4B). DFO-treated Mtb had significantly higher Rv1348 levels with insignificant changes in Rv1349 and Rv2895c levels (Figure 2F). DFO treatment for five days had a negligible impact on Rv2027c (*dosT*), Rv3132c (*dosS*), and Rv3133 (*dosR*) genes, which are known players in the induction of dormancy and persistor Mtb formation (Figure 2G).18 DAVID gene ontology clustering revealed >1.2–3.0-fold enrichment of transcripts encoding proteins having ATPase activity, indicating DFO’s impact on Mtb energy metabolism (Figure 2H). DFO also impacted the TCA cycle, oxidative phosphorylation, secondary metabolite synthesis, butanoate metabolism, fatty acid biosynthesis, and host-immune response (Figure 2H and I, Supplementary Figure S3B and C). DFO treatment led to upregulation of biological processes such as cellular response to starvation, chemical homeostasis, iron starvation, anaerobic respiration, and siderophore biosynthesis in Mtb (Figure 2I, shown in red in Supplementary Figure S3B). RNA modification, Heme metabolism, Nicotinamide, Cobalamin, Tetrapyrrole, Porphyrin, Arginine, Glutamine, Vitamins, Alpha-amino acid, Aromatic amino acid, branched-chain amino acid, aspartate biosynthesis/metabolism were downregulated upon DFO treatment (Figure 2I). Interestingly, downregulation of RNA modification, macromolecule methylation, fatty acid metabolism, dicarboxylic acid metabolic process and aromatic compound biosynthesis in DFO-treated Mtb were observed (shown in green in Supplementary Figure S3C). RNA-seq analysis of DFO-treated Mtb suggested that it has a direct impact on Mtb transcriptome and, as a result, may also alter the associated biological functions. A complete list of DEGs is included in Supplementary Table S1.

Earlier reports elegantly explained the role of various Mtb genes in iron metabolism and how iron uptake is a highly regulated process that can be disrupted for therapeutic intervention.^19^ Out of the reported genes, we observed upregulation of Rv3622c (PE32), Rv2353c (PPE39) and Rv3022A (PE29), and the rest showed marginal changes which could be due to experimental differences (Supplementary Table S2).

### DFO treatment alters Mtb metabolome

Further, the impact of DFO on Mtb metabolome was monitored using GC-MS analysis. PCA plot revealed that metabolites of DFO treated and control H37Rv Mtb clustered separately (Figure 3A and B). DFO-treated H37Rv Mtb showed significantly higher abundance (log2FC >± 1.0, -log10 p-value ≥ 1.3) of five metabolites (Citric acid, Ribofuranose, Erythrotetrofuranose, Myoinositol and Propanoic acid) compared to control (Figure 3C). Pathway impact analysis revealed that streptomycin biosynthesis, inositol phosphate metabolism and pentose phosphate pathway were significantly overrepresented in DFO-treated Mtb (Figure 3D). These findings indicate that DFO impacts the Mtb metabolome and causes detrimental effects on Mtb through iron deprivation. However, the target of DFO in Mtb is unknown, as with many antibiotics, including pyrazinamide. Next, we aimed to monitor whether DFO can limit intracellular Mtb growth and duplication.

**Figure 3:**
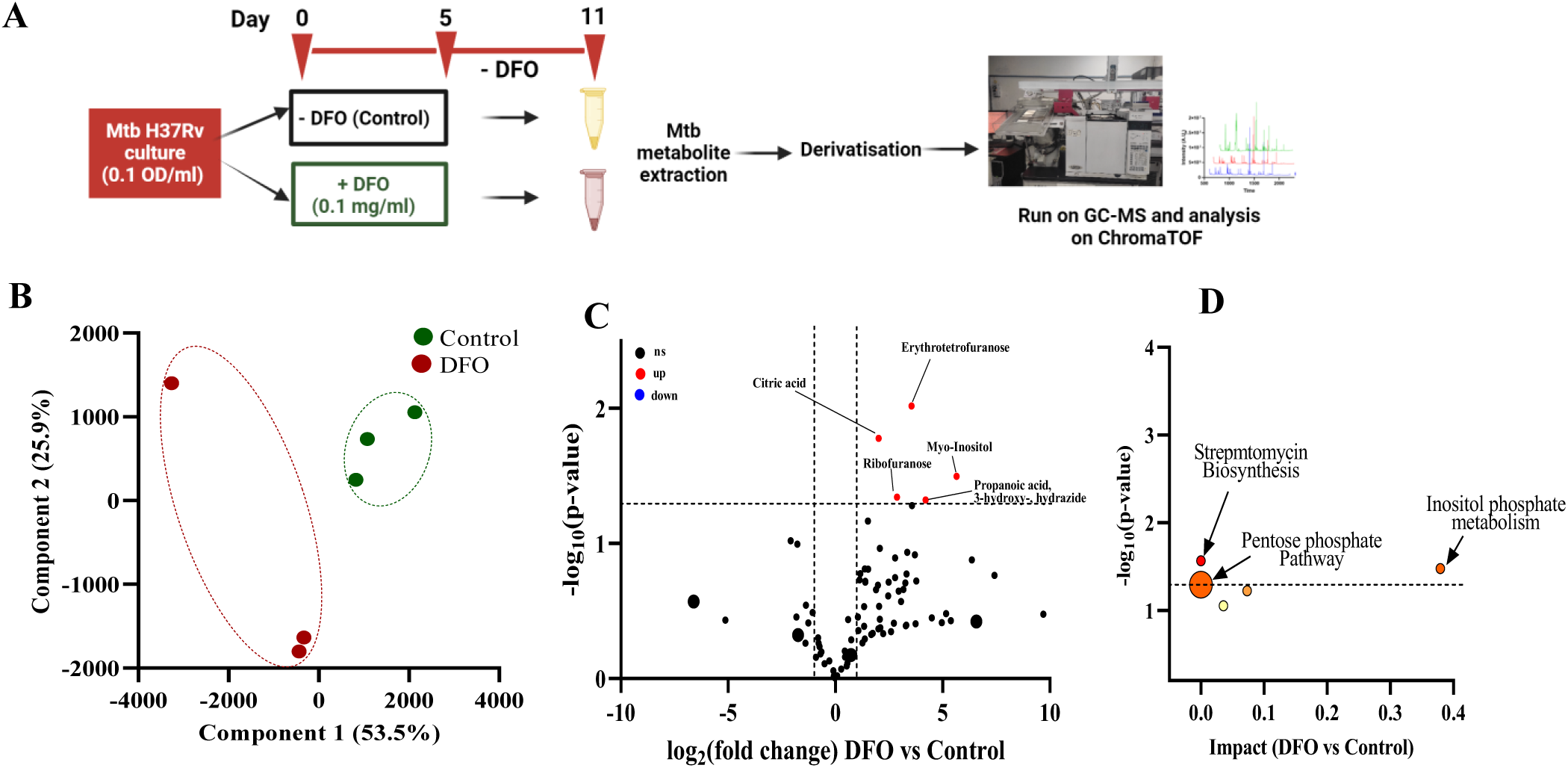
DFO alters Mtb global metabolite profile in-vitro. **A.** Schematic presentation of the method used for metabolite profiling from DFO-treated Mtb H37Rv and control using GC-MS. **B.** Principal component analysis showing DFO-treated Mtb H37Rv clustering separately from the control group. **C.** Deregulated metabolites observed between DFO-treated and control Mtb. **D.** Pathway analysis showing overrepresented metabolic pathways in DFO-treated Mtb.

### DFO shows anti-mycobacterial activity in Mtb-infected J774A.1 cell culture

The confocal images of the Mtb-infected J774A.1 cells that received DFO treatment or control were analysed using NIS-Element software. The DFO-pretreated J774A.1 cells showed decreased intracellular mycobacterial burden in a dose-dependent manner (Supplementary Figure S4C). Interestingly, we also observed hypertrophy, marked by increased cell size in DFO-treated cells compared to control.

### Iron supplementation leads to liver iron loading, higher tissue mycobacterial load and compromises the efficacy of INH and RIF

To validate the hypothesis that iron supplementation is not beneficial in TB patients, we created an iron overload mice model and infected it with a low aerosol dose of Mtb. The method adopted for iron overloading in C57BL/6 mice is presented in Supplementary Figure S5A. Prussian blue staining of the liver samples of iron-overloaded groups also showed higher iron accumulation and distribution (Supplementary Figure S5B, S7A and B). Iron overloading had a negligible effect on the mice’s body weight during and after the course (Supplementary Figure S5C). However, the iron-overloaded mice group showed significantly higher liver iron levels than the control group, and the INH-RIF treatment had a marginal reduction (Supplementary Figure S5D, S6A and B). Prussian blue staining and ICP-MS analysis of the liver of iron-overloaded mice showed increased iron levels till 75 days post-treatment (Supplementary Figure S6C, D and S7D, E). The tissue (lungs, spleen, and liver) mycobacterial burden was significantly higher in the iron-overloaded mice group at 15 days post-infection compared to the controls (Supplementary Figure S6E, F and G). The whole blood and thigh muscle iron levels of iron-overloaded mice did not show significant variations (Data not shown). As expected, the INH-RIF treatment cleared the tissue mycobacterial burden in control groups; however, the mycobacterial clearance in the lungs and spleen of iron overloaded mice groups was delayed and took longer time (>30 days of treatment) to bring it to lower than the limit of detection. Interestingly, the liver mycobacterial burden of the iron-overloaded group was similar to the control groups at 45- and 90-days post-infection. This demonstrated that higher host iron levels compromise the anti-mycobacterial efficacy of first-line anti-TB drugs.

### Iron chelation by DFO significantly reduces the tissue mycobacterial burden in C57BL/6 mice

Mtb infected C57BL/6 mice treated with DFO alone showed significant reduction in the liver iron levels as seen by Prussian blue staining and ICP-MS (Figure 4A, B, C and Supplementary Figure S7C, G). The decrease in iron levels was followed by a significant reduction in lung Mtb load within 30 days post-treatment and continued up to 75 days post-treatment compared to the infected control (90 dpi, Figure 4C and D). No change in the spleen mycobacterial burden was observed (Figure 4E). However, the liver of DFO-treated mice showed a significantly lower mycobacterial load (Figure 4F). This suggested that DFO alone has a direct effect that indirectly affects Mtb clearance in the lung tissues, so we studied the impact of DFO in combination with the anti-TB drugs in vivo in Mtb-infected mice.

**Figure 4:**
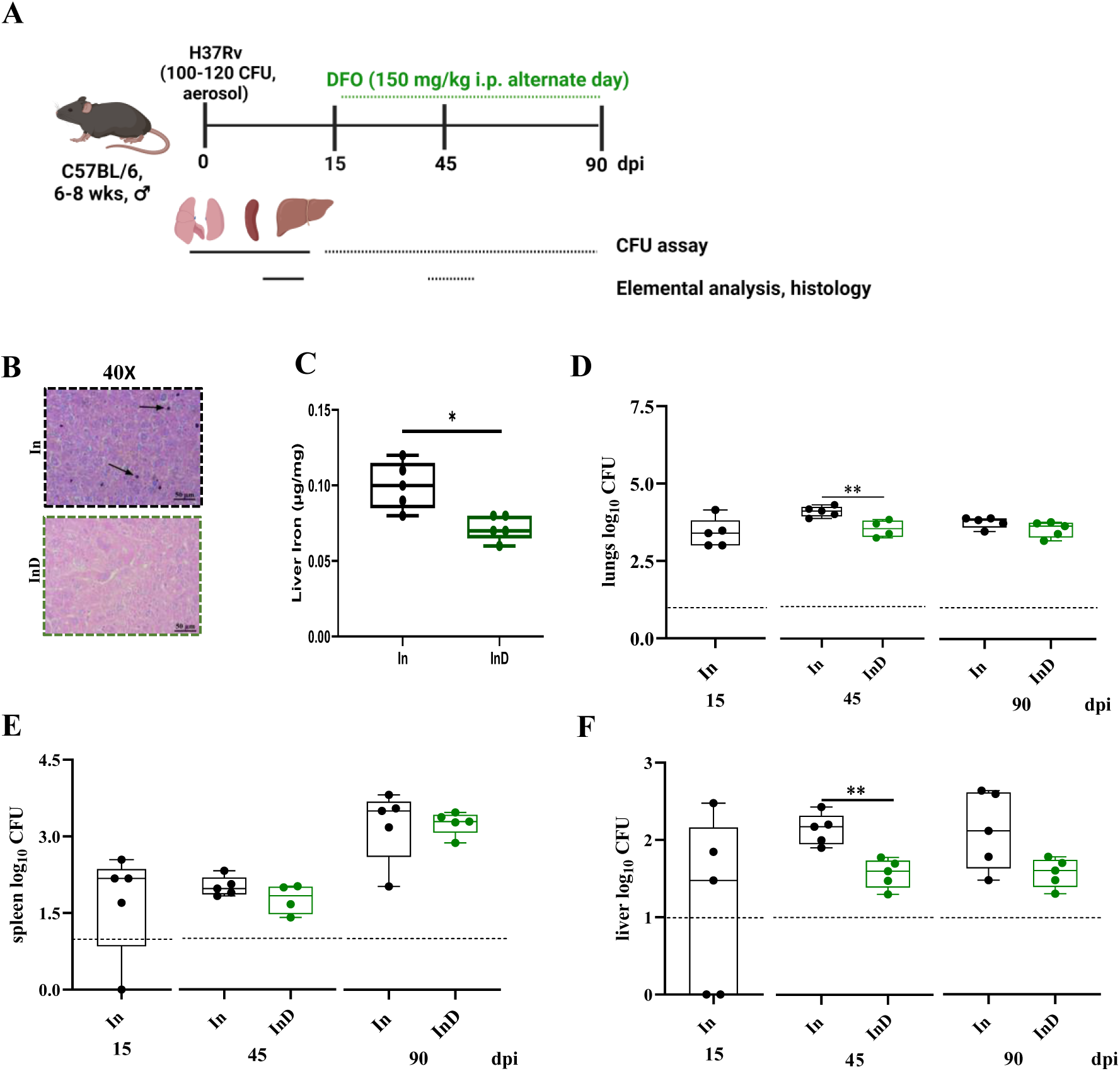
DFO aids in Mycobacterial clearance in C57BL/6 mice. **A.** Schematic presentation of the method used to infect C57BL/6 mice with *Mycobacterium tuberculosis* H37Rv strain and treatment using DFO. **B.** Prussian Blue stained liver histology for DFO-treated mice showing decreased liver iron accumulation indicated by black arrows. **C.** Liver iron levels were estimated using ICP-MS in the infected control compared with Mtb-infected mice receiving DFO treatment. Mycobacterial load in the lungs **(D)**, spleen **(E)** and liver **(F)** of the control and DFO treated *Mycobacteria tuberculosis* H37Rv infected C57BL/6 mice. Each time point had five biological replicates per group. The dashed line in the plots represents the LOD. i.p.: intra-peritoneal injection; dpi: days post infection; CFU: colony forming units. Results are determined using unpaired, non-parametric two-tailed t-tests. ns: not significant. *: p<0.05; **: p<0.01; ***: p<0.005; p-values at 95% confidence interval.

### DFO enhances lung bacterial clearance, increases recruitment of CD4^+^ T cells and macrophages impacting liver metabolite profile with the existing anti-TB drugs

Significantly lower liver iron levels were observed in RIF-INH treatment with or without DFO (Figure 5A, B, C and Supplementary Figure S7F, H). DFO adjunct to anti-tuberculosis drugs (InRHD) showed no difference in the lung bacterial burden at 30 days post-treatment (Figure 5D). Interestingly, Mtb-infected C57BL/6 mice treated with DFO adjunct to INH-RIF (grouped as InRHD) showed a significant reduction in lung mycobacterial burden at early time points, i.e., one- and two-weeks post-treatment, compared to the group receiving only INH-RIF (grouped as InRH) treatment (Figure 6A, B, and F). We monitored the immune cell distribution in an independent experiment and observed that InRHD group had a significantly higher frequency of CD4^+^ T cells and F4/80+ macrophages at one-week post-treatment compared to the InRH group and In and InRH groups, respectively (Figure 6B and C). The InRHD group had a significantly lower frequency of CD4^+^ T cells and F4/80+ macrophages at two-week post-treatment compared to the Mtb infected (In) group and group receiving TB drugs (InRH), respectively (Figure 6A and G).

**Figure 5:**
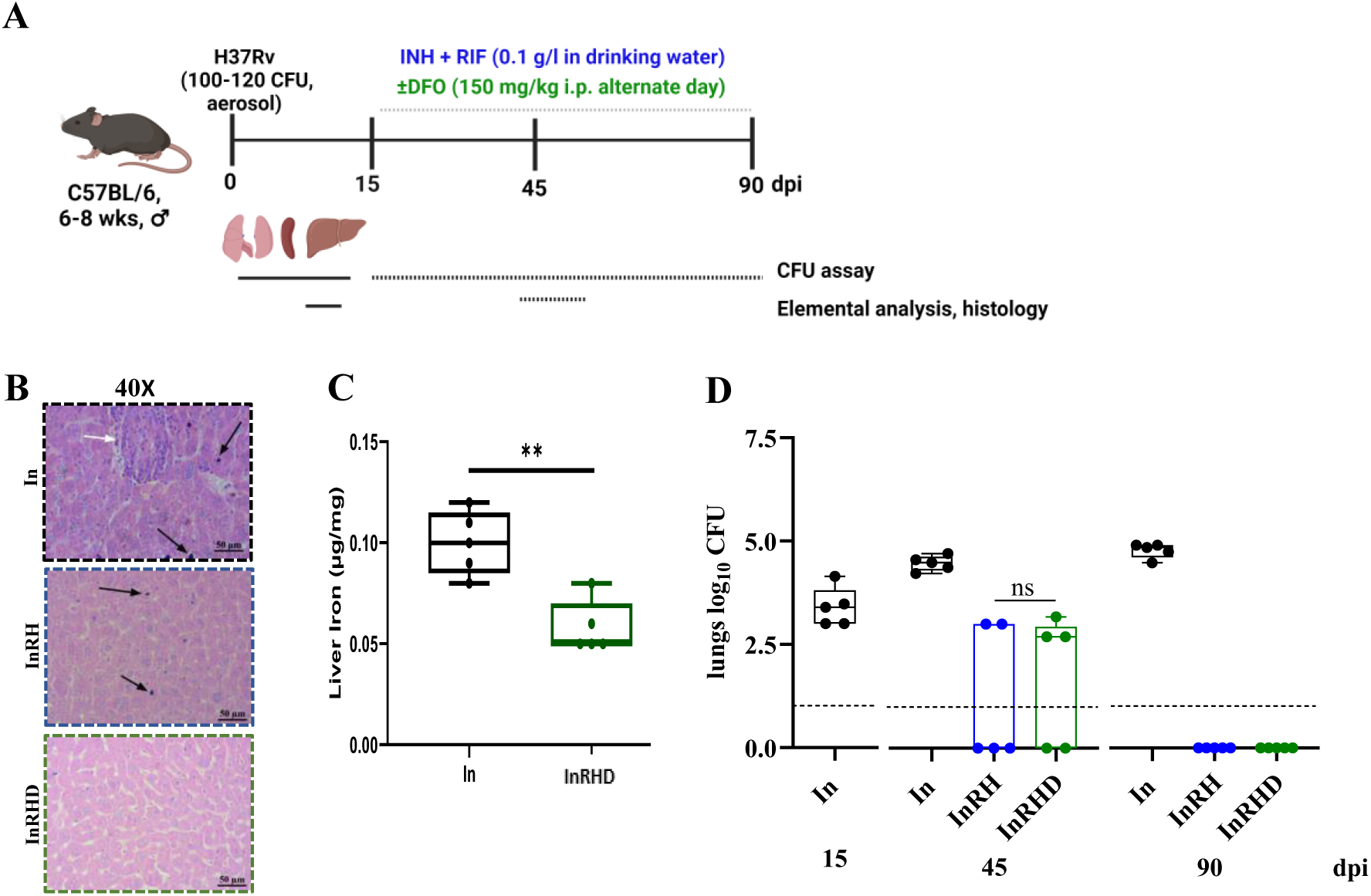
DFO adjunct to anti-tuberculosis drugs (INH/H and RIF/D) minimally affects lung mycobacterial burden at later timepoint. **A.** Schematic presentation of the method used for Mycobacterial infection and treatment using INH/H and RIF/R with or without DFO/D. **B.** Prussian blue staining of the liver tissue of RHD-treated mice (InRHD) shows decreased iron accumulation indicated by black arrows compared to IR-treated and infected control mice. **C** Liver iron levels estimated using ICP-MS in the infected controls and infected mice receiving INH and RIF with DFO. **D.** Lung mycobacterial load of the *Mycobacteria tuberculosis* H37Rv infected C57BL/6 mice receiving INH and RIF with or without DFO. Each time point had five biological replicates per group. The dashed line in the plots represents the LOD. i.p.: intra-peritoneal injection; dpi: days post-infection; wpt: weeks post-treatment, CFU: colony forming units. Results are determined using unpaired, non-parametric two-tailed t-tests. ns: not significant. **: p<0.01, ns: not significant; up: upregulated, down: downregulated, p-values at 95% confidence interval.

**Figure 6:**
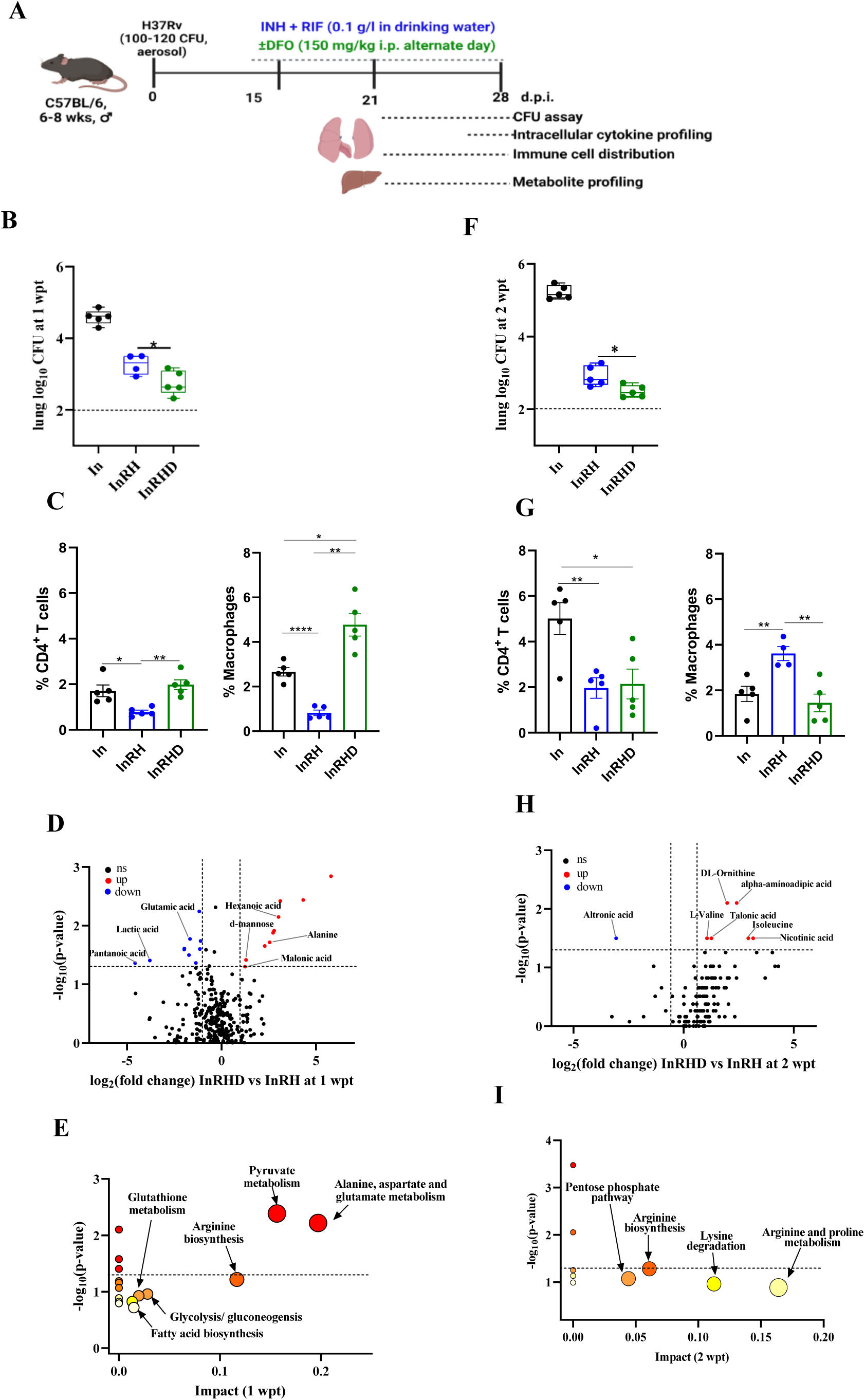
DFO adjunct to anti-tuberculosis drugs (INH/H and RIF/D) accelerated early mycobacterial clearance in C57BL/6 mice and impacts host liver metabolome. **A.** Schematic presentation of the method used for Mycobacterial infection, treatment using INH and RIF with or without DFO and monitoring host tissues at early time points **B.** Lung mycobacterial load one wpt. **C.** Frequency of CD4^+^ T cells and F4/80+ macrophages in the lungs of *Mycobacteria tuberculosis* H37Rv infected C57BL/6 mice receiving INH and RIF with or without DFO at the first week of treatment. **D.** Liver metabolome of Mycobacterial infected mice treated with anti-tuberculosis drugs with DFO vs without DFO at the first week of treatment. **E.** Scatter dot plot showing pathway enrichment analysis of deregulated metabolites from **D** plot. **F.** Lung mycobacterial load two wpt. **G.** Frequency of CD4^+^ T cells and F4/80+ macrophages in lungs of *Mycobacteria tuberculosis* H37Rv infected C57BL/6 mice receiving INH and RIF with or without DFO at the second week of treatment. **H.** Liver metabolome of Mycobacterial infected mice treated with anti-tuberculosis drugs with DFO vs without DFO at the second week of treatment. **I.** Scatter dot plot showing pathway enrichment analysis of deregulated metabolites from **H** plot. Each time point had five biological replicates per group except InRH group for immune cell distribution at 2 wpt. The dashed line in the plots represents the LOD. i.p.: intra-peritoneal injection; dpi: days post-infection; wpt: weeks post-treatment, CFU: colony forming units. Results are determined using unpaired, non-parametric two-tailed t-tests. ns: not significant. *: p<0.05; **: p<0.01; ****: p<0.0001, ns: not significant; up: upregulated, down: downregulated, p-values at 95% confidence interval.

Liver global metabolome analysis of InRHD vs InRH group completing the first week of treatment, using GC-MS, showed 20 deregulated (log2 Fold change >±1.0; p-value< 0.05) metabolites (Figure 6D). DFO treatment impacted the liver alanine, glutamic acid, lactic acid, and malonic acid levels (Figure 6D). Pathway enrichment analysis showed an overrepresentation of alanine, aspartate and glutamate, pyruvate metabolism and arginine biosynthesis in the treated group (Figure 6E). After one-week of INH-RIF treatment with or without DFO (both InRH and InRHD), liver pantenoic acid, lactic acid and alloxanoic acid levels were deregulated compared to the Mtb-infected control (In) (Supplementary Figure S8A and B). Similarly, seven deregulated metabolites were identified in the liver of the DFO-treated group, completing two weeks of treatment (Figure 6H). The liver of the InRHD group had significantly higher levels of proteogenic amino acids (valine and isoleucine) and non-proteinogenic amino acids (ornithine and aminoadipic acid) compared to the InRH group (Figure 6H). Arginine biosynthesis was overrepresented in the livers of mice receiving DFO as an adjunct to RIF and INH at 1 and 2 w.p.t. (Figure 6E and I). Long-chain fatty acids like oleic acid, arachidonic acid and octadecanoic acid were deregulated in the liver of the drug-treated mice (InRH, InRHD) as compared to the infected control (In) in the second week of treatment (Supplementary Figure S8C and D). The drug metabolism pathway for INH metabolism was enriched in the liver of mice groups receiving treatment (Supplementary Figure S9A, B and D). The arachidonic acid pathway was over-represented in the first week of treated groups (Supplementary Figure S9B). In contrast, biosynthesis of unsaturated fatty acids (linoleic and alpha-linoleic acid) was enriched in treated groups compared to the Mtb infected controls at first- and second-week post-treatment (Supplementary Figure S9C and D). These findings suggested that DFO accelerated Mtb clearance in C57BL/6 mice and impacted the liver metabolite levels significantly differently in adjunct to the INH and RIF than when treated with anti-TB drugs alone.

### DFO treatment adjunct to INH and RIF minimally effects cytokine production in the lung CD4+ T-cells

Increased CD4+ T cells and their proinflammatory cytokine releasing levels significantly improve lung Mtb clearance. Therefore, the effect of DFO on the frequency of CD4+ T cells and their pro-inflammatory cytokine production capacity after stimulation with PMA and ionomycin was monitored. The frequency of lung CD3^+^CD4^+^ cells in Mtb-infected C57BL/6 mice receiving INH-RIF with or without DFO at two weeks post-treatment was similar (Supplementary Figure S10A and B). The intracellular pro-inflammatory cytokines (IFN-γ, TNF-α and IL-17A) levels in the lung CD4^+^ T-cells of INH and RIF-treated groups with or without DFO were found to be similar (Supplementary Figure S10C). These findings suggested that DFO does not directly impact the CD4+ T cell frequency and pro-inflammatory cytokine production efficiency of CD4^+^ T cells, indicating that DFO does not hamper the effect of the immune system against active Mtb infection.

## Discussion

Numerous biological processes, including cell wall synthesis, ATP synthesis, DNA coiling, transcription, and translation in Mtb, have been instrumental in developing anti-TB drugs.^19^ However, limited studies have attempted to understand how limiting iron in the host affects Mtb growth and influences the effectiveness of anti-TB agents. Additionally, the effect of iron limitation on the survivability of Mtb has received less attention.^18,19^ In this study, we attempted to generate evidence of the pivotal role played by the host iron in Mtb survival and how manipulating iron availability through chemical agents, such as iron chelators, may impact Mtb clearance both *in vitro* and *in vivo* (Figure 7).

**Figure 7:**
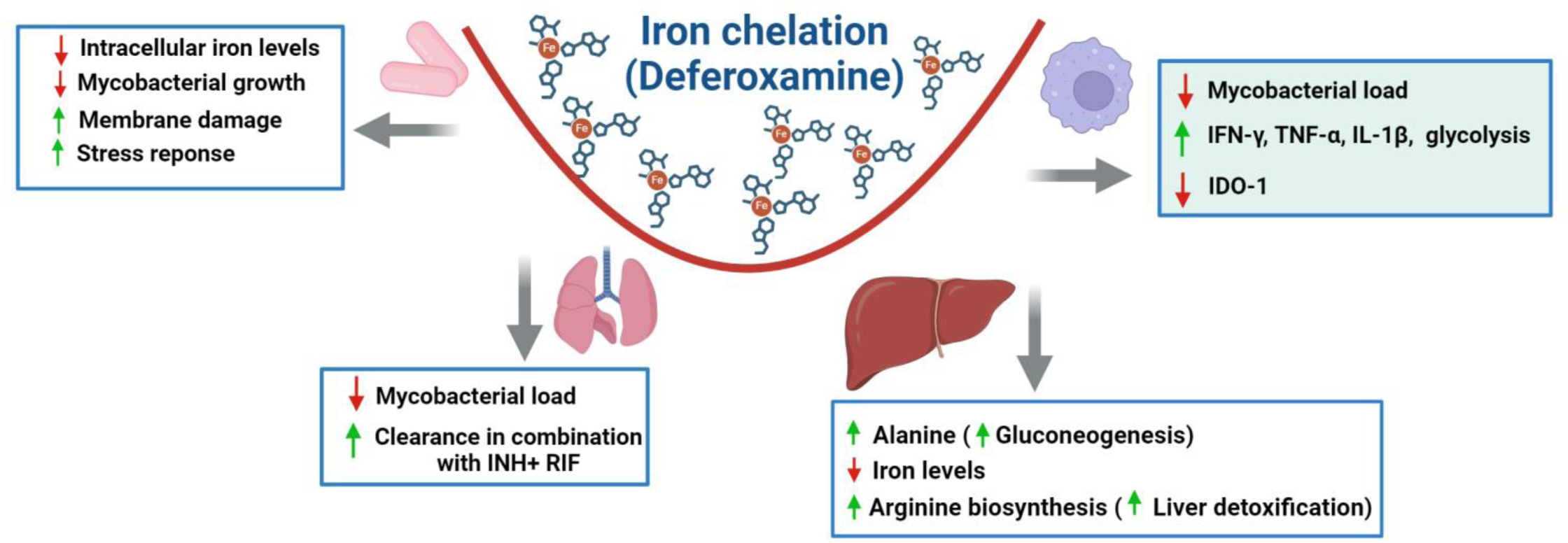
Schematic showing the effect of iron overloading and iron chelation on Mycobacterial growth, and tissues of Mtb infected mice. Mtb: Mycobacterium tuberculosis, INH: Isoniazid, RIF: Rifampicin, IDO: Indoleamine 2, 3-dioxygenase, TNF: Tumour necrosis factor, IFN: Interferon, IL: Interleukin.

Mtb cultured in 7H9 media with DFO exhibited notable bactericidal activity akin to the effectiveness of INH (Figure 1). Gallium nitrate is a ferric mimetic known to disrupt iron metabolism in Mycobacteria, and it decreases Mtb proliferation in infected macrophages.^18,19,20^ We observed that GN was less effective in inhibiting Mtb growth than DFO and INH *in vitro*. The observation of our *in vitro* study aligns with previous reports corroborating the anti-mycobacterial role of DFO.^9^ Iron deprivation targets mycobacterial membrane integrity by enhancing membrane permeability and disrupting membrane homeostasis reported in *M. smegmatis*.^25^ Electron micrograph images of H37Rv Mtb treated with DFO and INH showed cell wall damage, and a decreased envelope thickness compared to the controls (Figure 1F and Supplementary Figure S2). DFO also impacts Mtb’s intracellular copper and zinc levels (Supplementary Figures S1A and B). Transition metals like copper and zinc in low concentrations are essential for Mtb viability but, at high concentrations, can be toxic. Many enzymes in Mtb require these metal ions for their activity, like cytochrome C oxidase and superoxide dismutase, which require copper, copper, and zinc, respectively. So, the effect of DFO seems to be mainly due to iron deprivation and partly because of copper and zinc chelation as well.

Recent studies on iron deprivation in Mtb showed the up regulation of specific iron-containing proteins that aid in iron storage, genes involved in the uptake of iron and down regulation of the TCA cycle, and anabolic pathways for amino acid, vitamins, heme, and secondary metabolite biosynthesis.^16,17^ We also observed perturbed transcript levels of genes encoding proteins containing iron-sulfur clusters or iron alone. It seems that upon iron chelation by DFO, Mtb tries to scavenge the residual iron from surroundings by up regulating the genes involved in iron transport (Rv1348), iron-containing proteins (Rv1094, Rv1594) and storage (Rv0327c, Rv1553, Rv2276, Rv3251c, Rv3260c and Rv3406). RNA-seq analysis revealed that the gene signatures indicative of starvation and abiotic stress are affected, and DFO has a direct and detrimental effect on Mtb. Mtb is widely known to form a persister population when it encounters biotic and abiotic stress within the host or *in vitro*. However, the dos regulon encoded by Rv2027c(*dosS*), Rv3132c(*dosT*), and Rv3133c(*dosR*) did not show changes on day 5 of DFO treatment.18 Rv1348(irtA) seems critical for Mtb survival, as the CRISPRi knockdown assay revealed. The results suggested that DFO containing regimen may not lead to the generation of viable but non culturable bacteria (data not shown).

Mtb H37Rv global metabolite analysis revealed that DFO-induced metabolome changes persist even in the absence of DFO as myoinositol levels were significantly upregulated in DFO-pretreated Mtb as compared to Mtb cultured in rich media (7H9 supplemented with 10% OADC) (Figure 3C). Mycobacteria use myoinositol to synthesize phosphatidylinositol and mycothiol, an essential phospholipid required for cell wall synthesis and maintaining redox homeostasis in Mtb, respectively.^26^ Global metabolite analysis also indicated that DFO treatment affects the pentose phosphate pathway in Mtb. These results aligned with our transcriptomics analysis, which indicated that DFO treatment perturbs metabolic pathways in Mtb directly.

Several reports have shown that iron supplementation in mice, whether administered orally or peritoneally, supports the growth of non-tuberculous mycobacteria (NTM); however, the studies on Mtb are limited.^27^ Iron supplementation in mice significantly increased the Mtb burden in spleen and lung tissues aligning with our findings.^28^ Our study delved into the impact of host iron status on Mtb survival and growth *in vivo*, revealing that iron overloading achieved via intravenous administration of ferric carboxymaltose promoted higher mycobacterial load in the lung, liver, and spleen several weeks post supplementation (Supplementary Figure S8). However, this effect waned within a few months after discontinuing the iron overloading.

Limiting iron through DFO administration to Mtb-infected mice significantly reduced liver iron levels and mycobacterial burden compared to the infected control group (Figure 4C and 4F). Furthermore, liver iron levels in the iron-overloaded C57BL/6 mice significantly declined upon DFO treatment. We also assessed the effect of DFO when used as an adjunct to anti-TB drugs like INH and RIF. We observed substantially lower lung bacterial burden between animals treated with INH and RIF with and without DFO. It is critical to highlight that DFO administration, in conjunction with the existing TB drugs, reduced tissue Mtb load significantly at early time points. This suggests that DFO, as an adjunct to existing TB drugs, holds promise for potentially shortening the duration of anti-TB treatment by various mechanisms (Figure 7).

As the liver plays a critical role in regulating iron homeostasis by detecting even minor fluctuations in the systemic iron requirements and accumulating excess iron.^29^ Various regulatory mechanisms within the liver govern the expression of the iron regulatory genes, storage capacity, and iron mobilization. Dysregulation of these functions can lead to iron imbalance, a primary contributor to iron-related disorders. Iron supplementation in C57BL/6 mice had no discernible effect on their weight, corroborating earlier reports in Sprague-Dawley rat pups.^30^ To assess the effect of iron chelation on the liver, we observed lower liver mycobacterial burden in the DFO-treated groups. We also monitored the liver metabolome of Mtb-infected mice receiving INH and RIF treatment with or without DFO (Figure 6D and H). It is well known that anti-TB drugs cause drug-induced liver injury (DILI) in model organisms and humans. Global liver metabolite analysis showed an enriched arginine biosynthesis pathway in the DFO-treated group, which could be contributing to liver detoxification.^31^ Arginine, an essential antioxidant for the liver, is reported to neutralize ammonia, detoxifying the liver. Arginine supplementation helps in liver regeneration, and the production of antimicrobial compounds like nitric oxide (NO) in macrophages and as an adjunct to DOTS treatment, is beneficial to TB patients.^31,32^ Arginine, alanine, and valine abundance were higher in the liver of DFO-treated mice with potential immune-supportive functions.^33^ Alanine is converted to pyruvate by the alanine aminotransferase (ALT) enzyme in the liver, which contributes to glucose production by gluconeogenesis and enhances immune cell function against pathogens.^34^ DFO treatment enriched linoleic acid, alpha-linolenic acid metabolism, and arachidonic acid metabolism compared to Mtb-infected controls at the first and second week of treatment. Linoleic acid is also a precursor for arachidonic acid, which gives rise to an essential class of inflammatory mediators called eicosanoids to elicit an immune response against pathogens. DFO has been reported to increase glycolytic metabolism in Mtb-infected human monocyte-derived macrophages (MDM) while concurrently enhancing the early innate immune response by increasing the expression of IL-1β and TNF-α both at transcript and protein levels.^35^ Interestingly, we did not observe any significant changes in the intracellular cytokine levels (IFN-γ, IL-17A and TNF-α) in CD4^+^ T-cells harvested from the lungs of Mtb-infected C57BL/6 mice receiving DFO in combination with the first-line anti-TB drugs which may be a cell type-specific activity suggesting the direct role of DFO on macrophage cytokine production with minimal effect on cytokine production the CD4^+^ T-lymphocytes (Supplementary Figure S10C).^36^ But we did observe increased recruitment of CD4^+^ T cells in the lungs of the InRHD group at the first week of treatment but observed reduced frequency of these cells at the second week of treatment. It is likely that the lungs of mice receiving DFO adjunct to INH and RIF had lower CD4^+^ T cell and macrophage frequency because of effective bacterial clearance observed two weeks post-treatment. Importantly, DFO was observed to reversibly inhibit DNA synthesis in human B and T lymphocytes with limited impact on RNA and protein synthesis.^36^ In the presence of equimolar FeCl3, the effects of DFO in these cells were also reversed.^36^ We hypothesized that the combined effects of DFO with first-line anti-TB drugs could yield enhanced bactericidal effects and potentially reduce TB treatment duration. A recent report demonstrated that DFO treatment in BCG-infected primary hMDMs increased bedaquiline bactericidal activity, indicating its additive effects with the second-line anti-TB drugs.^9^ Further research is required to assess the efficacy of DFO in combination with these drugs against drug-resistant Mtb isolates. DFO can be introduced in the host by the subcutaneous, intramuscular, intravenous or oral route. Although iron chelation by DFO is the gold standard, it can also cause side effects like hypotension, fever, dysuria, leg cramps, rashes, pruritus, and free radical-mediated alveolar damage, leading to pulmonary toxicity, which intermittent infusions could avoid.

Targeting other critical trace elements like copper, zinc, and magnesium, which are essential for Mtb survival and growth, could also be a promising avenue for adjunct TB treatment strategies.^37,38,39^ Other iron chelators like Desferri-Exochelin may have a similar impact and need additional investigation.^40^ Additionally, exploring the role of gene polymorphisms in NRAMP1, the HFE gene associated with hemochromatosis, HP and α2 macroglobulin genes, which contribute to TB susceptibility, may facilitate the identification of the populations that could benefit the most from iron chelation therapy.^10,41^ Currently, there is no effective TB vaccine available for adults, and drug-resistant TB cases are on rise, it is crucial now than ever to invest resources in repurposing of FDA-approved or novel drugs for TB treatment. Several FDA-approved drugs have emerged as potential host-directed therapeutic candidates for combination therapy with the existing TB drugs. DFO, an FDA approved drug with pro-inflammatory properties for treating conditions like hemochromatosis in humans, which has shown the potential to bolster host immunity in the present study, leading to improved Mtb clearance and, therefore, has the potential to serve as an agent for host-directed adjunct therapy.^42^ Our findings on the earlier clearance of Mtb in DFO in combination to RIF and INH offers a new insights into manipulating host iron metabolism to shorten the treatment duration (Figure 7). Host-directed therapies involving iron chelation merit further validation to determine their impact on limiting Mtb drug resistance development, if any. Particular attention should be given to the potential toxicity of DFO treatment, and introducing an equivalent amount of FeCl_3_ in cellular systems negates its function.^30^ It is worth exploring whether DFO, as an adjunct, could also reduce the dosage of existing TB drugs, which presents an exciting avenue for future research. New therapeutic interventions targeting host or pathogen iron metabolism, such as iron chelation, are poised to contribute significantly to the fight against TB.

## Experimental procedures

### Ethical statement

The experiments performed in this study were approved by the Institute Biosafety Committee and the Institute Animal Ethics Committee (ICGEB/IAEC/07032020/TH-11) of the International Center for Genetic Engineering and Biotechnology, New Delhi.

### Mycobacterial H37Rv culture

*Mycobacterium tuberculosis* H37Rv strain (OD = 0.02) were inoculated in 7H9 media (BD Difco Middlebrook) supplemented with glycerol (0.2%), tween 80 (0.05%) and OADC (Oleate, albumin dextrose and catalase, 10%) in the absence or presence of individual drugs or in combination in the tuberculosis aerosol challenge facility (TACF, BSL-III) at ICGEB New Delhi. Mtb were grown at 37°C in a shaker incubator set at 180 rpm. Individual drug solutions were prepared at their minimum inhibitory concentrations (MICs) for DFO (0.1 mg/ml) and isoniazid (INH, 0.1 µg/ml) using milliQ water. Mtb growth was monitored by measuring absorbance at 600 nm at an interval of 24 hours up to 5 days post inoculation using a spectrophotometer to monitor the drugs’ bactericidal activity. On the fifth day, the harvested control and drug treated Mtb were inoculated on 7H11 agar plates supplemented with OADC (10%) and BD MGIT™ - PANTA™ antibiotic Mixture for colony forming unit (CFU) assay. On the fifth day, a fraction of the Mtb culture (0.8 OD, 133 M) were centrifuged at 3,500 g for 10 minutes at room temperature and elemental analysis was performed in the pellet and the cultured media as a separate fraction by using inductively coupled plasma mass spectrometry (ICP-MS) analysis (Figure 1A).

### Transmission electron microscopy of Mtb

Mtb cultures (0.1 OD) were inoculated with DFO, INH, and Gallium nitrate (GN, 4µg/ml) at MIC and control, incubated at 180 rpm at 37°C. Mtb cultures were harvested 5 days post-treatment and centrifuged at 3,500 g for 5 minutes, followed by washing with phosphate buffer saline (PBS) by centrifuging at 3,500 g for 5 minutes. The pellet containing Mtb was fixed with 2% glutaraldehyde for an hour. The fixed bacterial pellet was washed twice with PBS and then passed five times through a 26 G needle to make a single-cell suspension resuspended in 100 µl ddH_2_O. 10 µl suspension was loaded onto a carbon grid and stained with 5% uranyl acetate. After air drying the grid, it was loaded in the Transmission Electron Microscope (Tecnai G2 spirit, HT120kV) and the micrographs were captured (Figure 1E).

### Kill kinetics of Mtb in the presence of DFO, GN, INH and in combination

Log phase (OD= 0.2) Mtb H37Rv culture were inoculated in 7H9 media (2 ml) supplemented with OADC (10 %), glycerol (0.2 %) and tween 80 (0.05 %) in presence of 10 × MIC of DFO (1 mg/ml), GN (40 µg/ml), INH (1 µg/ml) and in combination (DIG: DFO +INH + GN) and were incubated at 37°C at 180 rpm. Aliquots (100 µl) of each culture condition were harvested at days 0, 4 and 8 for CFU assay by plating diluted samples on 7H11 agar plates supplemented with OADC (10 %) and BD MGIT™ - PANTA™ and incubated at 37°C. Mycobacterial colonies were enumerated after 3 weeks post-plating.

### Transcriptome analysis of DFO treated Mtb

Mtb H37Rv cultures (0.1 OD/ml) treated with DFO (0.1 mg/ml) and control were harvested on day 5 by centrifuging at 3,500 g for 10 minutes at room temperature. The pellet after washing with Tris-EDTA (1 M) to remove extra media and resuspended in Trizol reagent (1 ml, Invitrogen Inc., USA). The samples were subjected to bead beating (4 cycles, 45 sec each) after adding zirconia beads (∼200 μl, 0.1 mm) at 6.5 m/s speed with incubation of 30 sec on ice after each cycle using Biospec Mini-Beadbeater-16. Chloroform (0.2 ml) was added to the samples and after briefly shaking the tubes, the samples were incubated on ice (for 5 min) and centrifuged at 13,400 g, 4°C for 20 min. The upper aqueous layer was transferred to a fresh tube and RNA was precipitated by adding isopropyl alcohol (0.5 ml). After incubating the samples at -80°C for 20 min, were centrifuged at 13,400 g for 10 min at 4°C. The pellet was washed with 75% ethanol and resuspended in nuclease-free water. RNA was quantified using a Qubit BR RNA estimation kit (Invitrogen Inc., USA) as per the manufacturer’s protocol and RNA integrity was assessed using Agilent TapeStation (Agilent Inc., USA). Samples (0.5 μg each) above a RIN value of 7 were sent for RNA sequencing to NGB Diagnostics Pvt. Ltd. Briefly, library preparation was performed using NEBNext Ultra RNA library prep kit (NEB, USA). The uniquely indexed libraries were sequenced on the Illumina NovaSeq 6000 platform with 150×2 paired-end chemistry (Figure 2A).

The Next Generation Sequencing (NGS) data in fastq format was analyzed using FastQC (v0.12.1) to determine the quality and adapter contamination. Adapter contamination was addressed using bbduk.sh (from BBmap toolkit v39.06) with options (ktrim=l k=23 mink=11 hdist=1 tpe tbo). Hisat2 (v2.2.1) was used for the alignment of reads to the Mtb genome (Refseq Accession: NC_000962.3) which was subsequently sorted using Samtools (v1.13). The sorted alignment files in BAM format were used as input for counting reads using featureCounts (v2.0.3). The read count matrix was formatted so that it can be used as input for iGEAK! and iDEP.96. EdgeR was used to identify the differentially expressed genes with a minimum CPM value of 1, a log_2_DFO/control>1.0 and adjusted p-value ≤ 0.05 calculated using Benjamin-Hochberg method. To identify the deregulated pathways, the same raw read matrix was analyzed using the pre-ranked FGSEA option in iDEP.96.

### CRISPRi knockdown strain preparartion

The pLJR965 CRISPRi plasmid (kind gift by Dr. Sarah M. Fortune) backbone was digested with BsmBI. Subsequently, sgRNAs designed to target the *irtA* ORF was cloned using previously established methods.^40^ Confirmation of successful cloning was obtained through partial sequencing. The resultant plasmid containing the *irtA*-targeting sgRNA was then introduced into Mtb via electroporation. Bacterial cultures were allowed to recover in 7H9 medium for 24 hours, after which were plated onto 7H11 agar supplemented with kanamycin (50 µg/ml) for the selection of transformants. The inducible system was triggered by anhydrotetracycline (Atc, 50 ng/ml) supplementation on every third day.

### Global metabolite analysis of DFO-treated Mtb H37Rv

Mtb H37Rv cultures (normalized with bacterial number) treated with DFO (0.1 mg/ml) and control were harvested on day 5 by centrifuging at 3,500 g for 10 minutes at room temperature (Figure 3A). Mtb pellet harvested from DFO-treated and control condition was incubated in drug-free 7H9 media with OADC (10%) for 6 days and then equal number of bacteria were harvested for metabolomics analysis. The pellet was resuspended in 80% methanol along with ribitol (2 μl, 0.5 mg/ml) as a spike in standard and transferred to bead beating tubes containing 0.1 mm zirconia beads. The samples were subjected to bead beating using Biospec Mini-Beadbeater-16 at 3650 oscillations/minute, 30 seconds on/off cycles six times with incubation on ice during the off cycle. After bead beating, the samples were incubated on ice for 30 minutes followed by centrifugation at 10,000 g for 10 minutes at 4°C to collect the supernatant (800 μl). The extracted metabolites were filtered using a 0.2 μm, nylon membrane filter (#726-2520) and stored at -20°C until further processing. Equal volume of the extracted metabolites from all the samples were pooled to prepare two QC samples.

For derivatization, samples were dried at 40°C using vacuum concentrator (Labconco Centrivap, USA) and incubated with Methoxamine hydrochloride (MeOX-HCl, 2%, 40 μl) at 60°C for 2 hours at 900 rpm in a thermomixer (Eppendorf, USA). After adding N-methyl-N-(trimethylsilyl) trifluoroacetamide (MSTFA, 70 μl), it was incubated at 60°C for 30 minutes at 900 rpm in a thermomixer. After incubation, the samples were centrifuged at 10,000 g for 10 minutes at 25°C, the supernatant was transferred to GC vial inserts for GC-MS analysis using a 7890 Gas Chromatography (Agilent Technologies, Santa Clara, CA) coupled to a Pegasus 4D GC × GC time-of-flight mass spectrometer (LECO Corporation). The samples were randomized using Research Randomizer (www.randomizer.org) and processed together with one QC sample ran before and after the sequence. The derivatized samples (1 μL) were injected to an HP-5ms GC column (30 meters length, 250 µm internal diameter) in splitless mode using Helium as a carrier gas at a constant flow rate (1 ml/min) and secondary column was Rxi-17 of 1.5 m length and 250 µm diameter. Electron ionization (EI) mode was fixed at −70 eV to scan ions of 33 to 600 m/z range at acquisition rate of 20 spectra/second. The ion source temperature was set at 220°C. GC method parameters used for acquisition were: 50°C for 1 min, ramp of 8.5°C/min to 200°C, ramp of 6°C/ min to 280°C, hold for 5 minutes. Secondary oven temperature offset was set at +5°C relative to GC oven temperature. Similarly, the modulator temperature offset was set at +15°C relative to the secondary oven temperature. Transfer line temperature was set at 225°C. A solvent delay of 600 seconds was used, and all data acquisition was completed within 24 hours of derivatization. A commercial standard mix of amino acids were derivatized and run using the same GC-MS method to confirm the identity of the important metabolic features. All GC-MS raw data files of the study groups were aligned using the “Statistical Compare” feature of ChromaTOF (4.50.8.0, Leco, CA, USA). Peaks with a width of 1.3 seconds and the signal to noise ratio (S/N) threshold of ≥50 were selected. Putative identity of the peaks was assigned using a NIST (National Institute of Standards and Technology, USA) library (version 11.0; 243,893 spectra) search qualifying a similarity match of >600 at a mass threshold of 100. The .csv file, generated from the above analysis, was manually curated to check the peak identity. Analytes present in more than 50% of the samples of a study group were selected for statistical analysis. Ribitol peak area from the sample was used for normalizing the liver metabolite data. Fold change analysis and p-value were calculated using MetaboAnalyst software (version 6.0). Metabolic features that showed log_2_ fold change ≥ ±1.0 with p-value < 0.05 between the study groups were selected as deregulated molecules. Pathway enrichment analysis for deregulated molecules was performed using Pathway analysis plugin in the MetaboAnalyst software.

### J774A.1 infection with H37Rv and DFO treatment

The J774A.1 cell was seeded on sterile round cover glass in a 6-well plate. The cells were pretreated with different DFO concentrations (0, 0.25, 2.5 and 25 mM) for 24 hours before introducing Mtb H37Rv at 1:10 multiple of infection (MOI). After washing, the cells were stained with DAPI (1 μg/ml, Invitrogen #D1306), MitoTracker™ Red CMXRos (200 nM, Invitrogen #M7512) and fixed at different time points and finally slides were prepared for confocal microscopy by using the prolong gold antifade mountant (Invitrogen, Cat# P36934). These fixed cells were used for fluorescence microscopy using a laser scanning confocal microscope (Nikon A1R, USA), and images were analyzed using NIS-Element software.

### Iron supplementation in C57BL/6 mice and Mtb aerosol infection

C57BL/6 mice (8-10 weeks old, male) from the host institute (ICGEB) animal house were transferred to the tuberculosis aerosol challenge BSL-III facility (TACF) for adaptation for at least one week prior to the infection experiment. The animals were kept in a specific pathogen-free environment, 12-hours daylight conditions, and food and water were provided *ad-libitum*. A set of the study mice received ferric-carboxymaltose (Ferinject, LUPIN ltd, India, 0.5 mg iron/mice in 100 µl saline) through intravenous injection, while the control mice received saline (100 µl). These iron-supplemented mice received a total of four doses of ferric-carboxymaltose injections on every fourth day. After a week of rest, the iron-supplemented and control mice were aerosol infected with a low dose (100-120 CFU) of Mtb H37Rv strain using Madison chamber present in TACF and sacrificed at 15-, 45- and 90-days post-infection (dpi) to monitor the tissue bacterial load. A subgroup of Mtb infected C57BL/6 mice received either INH-RIF (InIR, 0.1 g/l each, changed alternate day) in drinking water ad-libitum or DFO (InD, 150 mg/kg body weight of mice, intraperitoneal injection) or both (InIRD) starting 15 dpi for 75 days (i.e., 90 dpi). DFO dose was introduced intraperitoneally every alternate day till treatment completion. At the end of the study, mice were anaesthetized using 5% Isoflurane, and organs were harvested from the humanely euthanized mice. Harvested tissues (lungs, spleen, and liver) were processed using a homogenizer in sterile PBS (1 ml) and an aliquot (100 µl) of it was inoculated on 7H11 agar plates supplemented with OADC (10%) and BD MGIT™ - PANTA™ antibiotic mixture at appropriate dilution in sterile PBS and the colonies were counted post 3 weeks of incubation at 37°C in the incubator. To a part of the liver tissue, chilled methanol was added and stored at -80°C for elemental analysis using ICP-MS and metabolite profiling using gas chromatography-mass spectrometry (GC-MS).

### Tissue histology analysis

A part of the harvested tissues was stored in 10% formaldehyde for histology analysis. Briefly, processed tissues were paraffin-embedded and 5 µm thick sections were used for slide preparation at an accredited pathology lab. Prussian blue staining was used for monitoring the tissue iron distribution. Histology slides were scanned in Zeiss microscope and analyzed using Zen 3.1 software (Supplementary Figure S7).

### Elemental analysis using ICP-MS

Equal number of bacteria (0.8 OD) were pelleted and to the pellet, HNO_3_ (70%, 150 µl) was added and transferred to sample vials (MG5, Anton Paar, USA) and then H_2_O_2_ (30%, 50 µl) was added. Similarly, to 1 ml culture filtrates, HNO_3_ (70%, 500 µl) and H_2_O_2_ (30%, 150 µl) were added. The sealed sample vials were subjected to microwave (Anton Paar, USA) digestion using a ramp of 250W for 15 min with a hold time of 5 min at 250W. Similarly, liver , thigh muscle (50 mg each) tissues or equal volumes of whole blood (50 µl) were taken in a digestion vial (64MG5 vials) and after adding HNO_3_ (400 µl) and H_2_O_2_ (100 µl) were incubated for 20 min at room temperature, and then the vials were sealed and subjected to microwave digestion. The initial power was set at a ramp of 10 min and 150W with a max temperature of 140°C, followed by a 15 min hold. Then, a second ramp of 15 min from 150-250W was used, and a final hold was used to bring the temperature to 55°C at the highest fan speed. After diluting the samples with trace metal-free water, iron (57Fe) levels were monitored in the digested samples using ICP-MS (iCAPTM TQ ICP-MS, Thermo Scientific, USA). Thermo Scientific Qtegra^TM^ Intelligent Scientific Data Solution (ISDS) software was used to operate and control the instrument. Briefly, the torch was warmed up for 30 min in single-quad Kinetic energy discrimination (SQ-KED) mode with helium to remove the polyatomic nuclei interference and then autotuned in normal mode followed by advanced KED mode. Then, the multi-element standard (#92091, Sigma Aldrich, USA) of different concentrations was run at the same settings to prepare the standard plot. A sample blank was run before each sample, and a quality control (QC) of known concentrations of standards was run after every 15–20 samples to monitor and maintain uniform signal intensity throughout the run. Data analysis was done using Qtegra software (Thermo Scientific, USA).

### Flow sorting of CD4^+^ T cells and macrophages from the lungs

We performed an independent experiment to monitor the immune cell distribution in the lungs of Mtb infected C57BL/6 mice. The right lobes of the lungs were harvested and incubated with collagenase (2 mg/ml) and DNase (1 mg/ml) in RPMI-1640 (2 ml) at 37°C for 30 min. The enzymatically treated tissues were passed through a strainer (70 µm) before centrifuging at 400 g for 10 min to collect the pellet. Ammonium chloride potassium (ACK) lysis buffer was added to the pellet, incubated for 2 min at room temperature and quenched by adding RPMI-1640. The cells were centrifuged and resuspended in RPMI-1640 and counted using a hemocytometer.

10 million cells from each sample were incubated with an antibody cocktail of LIVE/DEAD, CD45, CD3, CD4 and F4/80 surface markers (Live dead fixable stain #L34975, CD4-SB600 #63-0042-82 from Invitrogen and CD3-Alexa Fluor 700 #152316, CD45-PerCP/Cy5.5 #103132 and F4/80-PE #123110 from BioLegend) for 40 mins on ice. After staining, these cells were fixed using paraformaldehyde (4%) washed with FACS buffer and finally resuspended in the FACS buffer. Immune cell sorting was done using BD FACS Aria III Fusion (BD Biosciences, USA) present in TACF, and the data was analyzed using FCSExpress (DeNovo software, Version 6.0).

### Liver metabolite profiling using GC-TOF MS

For metabolites isolation, liver tissues (100 mg) were finely chopped and transferred to bead-beating tubes (2 ml) with zirconium beads (2 mm, 250 mg). Chilled methanol (80%, 1 ml) was added to the tubes along with ribitol (5 μl, 2 mg/ml) as a spike in standard. The tissues were subjected to bead beating using Biospec Mini-Beadbeater-16 at 3650 oscillations/minute, 30 seconds on/off cycles six times with incubation on ice during the off cycle. After bead beating, the samples were incubated on ice for 30 minutes followed by centrifugation at 10,000 g for 10 minutes at 4°C to collect the supernatant (800 μl). The extracted metabolites were filtered (0.2 μm, nylon membrane filter #726-2520) and stored at -20°C until further processing. Equal volume (120 μl) of the extracted metabolites from all the samples from a particular time point were pooled to prepare a QC sample. For liver metabolite derivatization, data acquisition and analysis using GC-MS, the same procedure as followed for global Mtb H37Rv metabolite analysis was adopted.

### Intracellular cytokine staining and flow cytometry

The right lobes of the lungs were harvested and incubated with collagenase (2 mg/ml) and DNase (1 mg/ml) in RPMI-1640 (2 ml) at 37°C for 30 min. The enzymatically treated tissues were passed through a strainer (70 µm) before centrifuging at 400 g for 10 min to collect the pellet. ACK lysis buffer was added to the pellet, incubated for 2 min at room temperature and quenched by adding RPMI-1640.

The cells were centrifuged and resuspended in RPMI-1640, then counted before seeding in a 12-well plate and incubated overnight with phorbol 12-myristate 13-acetate (5 ng, PMA) and ionomycin (0.75 µg). After washing, these cells were incubated with brefeldin (5 µg) and monensin (1.3 µg) for 3 hrs. The cells were washed with FACS buffer (1% FBS in PBS) and incubated with an antibody cocktail of LIVE/DEAD, CD3 and CD4 surface markers (Live dead fixable stain #L34975, CD3-FITC #11-0032-82 from Invitrogen, USA and CD4-BV785 #100552 from BioLegend) for 40 mins on ice. After staining, these cells were fixed using paraformaldehyde (4%) and permeabilized using Permeabilization buffer (#00-8333-56, Invitrogen, USA). These cells were incubated with an antibody cocktail of intracellular cytokines: TNF-α, IL-17A and IFN-γ (TNF-α-APC #17-7321-82, IL-17A-efluor450 #48-7177-82, IFN-γ-PerCP Cy5.5 #45-7311-82 from Invitrogen, USA). Flow cytometry data were acquired using BD LSR Fortessa X-20 (Supplementary Figure S10A) and analyzed using FCSExpress (DeNovo software, Version 6.0).

### Statistical analysis

All data are presented as mean ± standard deviation (SD) values. Statistical analyses were performed using Origin 2020, GraphPad Prism (Version 8.4.2) and Microsoft Excel using an unpaired two-tailed t-test to identify group-specific variations. Differences were considered statistically significant with p<0.05 at 95% confidence.

### Data availability

All the related data is presented in the manuscript and the transcriptome data generated from DFO-treated Mtb, and control could be accessed from NCBI-Gene Expression Omnibus (GEO) using accession ID: GSE264613.

## Supporting information

This article contains supporting information.

## Acknowledgements

We acknowledge the Department of Biotechnology (DBT), Government of India, for supporting activities through research grants to RKN and supporting the Tuberculosis Aerosol Challenge Facility at ICGEB and ICGEB New Delhi for providing access to the ICP-MS facility and core support to RKN. Sandeep R. Kaushik received a Senior Research Fellowship, and Nidhi Yadav and Ashish Gupta received a Junior Research Fellowship from the DBT. Sukanya Sahu received a Senior Research Fellowship, and Poonam Dagar received a Junior Research Fellowship from the Council of Scientific and Industrial Research (CSIR) in New Delhi. Nikhil Bhalla is supported by the DBT-funded project (Grant ID: National Network Project of National Institute of Immunology, New Delhi -[40267]). We thank Ms Mothe Sravya for helping with manuscript preparation.

## Author contributions

SRK, NY, and RKN conceptualized and designed experiments; SRK, SS, AKM, N and AG performed the animal experiments; SRK, SS, AKM, NY, AG, PD and AS carried out *in vitro,* and the laboratory profiling experiments; NB analyzed the Mtb RNA sequencing data; SRK and NY analyzed the data and prepared the figures; TS and AKP generated the CRISPRi knockdown of the Mtb strain; BB shared resources and reagents to carry out the project; RKN generated funds for this work; SRK and RKN wrote the first draft of the manuscript and revised it, incorporating the comments of all co-authors.

## Conflict of interest

The authors declare that they have no conflicts of interest with the contents of this article.

## Supplementary Table and Figure legends

**Supplementary Figure S1: A and B.** Intracellular copper and zinc levels in drug-treated and control Mycobacterial culture at 5th-day post-treatment quantified using ICP-MS. Results are determined using unpaired, non-parametric two-tailed t-tests. n = 5 per group, ns: not significant; *: p<0.05, **: p<0.01; ****: p<0.0001 at 95% confidence interval.

**Supplementary Figure S2:** Transmission electron micrographs of untreated Mtb **(A)**, DFO treated **(B)**, INH treated **(C)**, GN treated **(D)** harvested at 5th day post-drug treatment. Images with dashed boundaries are referred in the main figures.

**Supplementary Figure S3: DFO alters iron metabolism and various pathways of Mtb H37Rv. A**. Heatmap showing the deregulated (up in red and down in green) genes in the DFO (D1, D2, D3) vs control Mtb (C1, C2) as evident from the Z-score values. **B.** Pathway analysis using Preranked FGSEA in IDEP.96 tool. Red indicates upregulated and green shows down-regulated pathways in Mtb. The intensity of the colour and diameter of the node corresponds to the number of genes impacted in a particular pathway. **C.** Iron-sulfur cluster and iron-binding proteins impacted upon DFO treatment. Green shows presence and Black shows absence of the Iron-sulfur/Iron-binding nature of the protein.

**Supplementary Figure S4: irtA knock down in Mtb inhibits its growth and DFO exposure reduces Mtb growth in J774A1 cells. A.** Relative suppression of *irtA* and *irtB* expression levels in *irtAKD* and cntrl sgRNA strains (mean ± SEM, n = 3 biological replicates). **B.** Relative suppression of *irtA* and *irtB* expression levels in *irtAKD* and cntrl sgRNA strains **C.** Fluorescent confocal microcopy images acquired at 60X magnification showing the effect of DFO pretreatment on Mtb H37Rv infected J774A.1 murine cells. Blue color represents nuclei stained with DAPI and red color indicates mitochondria stained with MitoTracker red. Results are determined using unpaired, non-parametric two-tailed t-tests **P ≤ 0.005 and ***P ≤ 0.0005, ****P 852 ≤ 0.00005. Data represents mean ± SEM (standard error mean). (KD=knock down, cntrl=control)

**Supplementary Figure S5:** A. Schematic representation of the method used to create iron loading conditions B. Prussian blue staining of the liver tissue at 40X magnification indicating high iron accumulation shown by black arrows in the FcHealthy C57BL/6 mice as compared to healthy controls C. Body weight of the C57BL/6 mice before, during and after iron loading for FcHealthy in comparison to healthy controls D. Liver iron levels, estimated using ICP-MS at different time points during iron loading as compared to healthy controls. Scale bar for histology images: 50 μm, Fc: ferric carboxymaltose supplemented group. Results are determined using unpaired, non-parametric two-tailed t-tests where ns: not significant; **: p<0.01; ***: p<0.001 at 95% confidence interval.

**Supplementary Figure S6:** A. Schematic presentation of the method used to create iron loading condition, Mycobacterial infection and treatment (with isoniazid: INH and rifampicin: RIF) experiment. B. Liver iron levels, as estimated using ICP-MS, in iron loaded and control mice groups. C. Prussian Staining of liver tissues at 40X magnification indicates high iron accumulation shown by black arrows in iron-loaded mice and did not change significantly upon drug treatment (D). Mycobacterial load in the lungs (E), spleen (F) and liver (G) of the Mycobacteria tuberculosis H37Rv infected C57BL/6 mice. Dashed lines in the CFU plots represent the limit of detection. n = 5/group/time points; i.v.: intra-venous injection; dpi: days post-infection; CFU: colony forming units; scale bar for histology images: 50 μm. Results are determined using unpaired, non-parametric two-tailed t-tests where ns: not significant, p values at 95% confidence interval.

**Supplementary Figure S7:** Histology images of liver tissue stained with Prussian blue staining showing iron accumulation in H (A), FcH (B), In (C), FcIn (D), InRH (E), FcInRH (F) as compared to minimum iron in InD (G) and InRHD (H) groups at 4X, 10X, 40X magnification respectively. Scale bar: 50μm, In: Mtb infected C57BL/6 mice; FcIn: Mtb infected Fe supplemented C57BL/6 mice; InD: Deferoxamine (DFO) treated Mtb infected mice; InRH: Isoniazid (INH) and Rifampicin (RIF) treated Mtb infected C57BL/6 mice; FcInRH: INH and RIF treated Mtb infected iron supplemented C57BL/6 mice; InRHD: Mtb infected C57BL/6 mice receiving both DFO, INH and RIF; Healthy: Control mice (H); FcHealthy: Control mice with iron supplementation (FcH).

**Supplementary Figure S8:** Liver metabolome changes upon treatment with INH and RIF with or without DFO at 1st and 2nd weeks. InRH vs In **(A)** and InRHD vs In **(B)** InRH vs In **(A)** and InRHD vs In. ns: non-significant, up: upregulated, down: down regulated, p value at 95 % confidence interval, In: H37Rv infected control, InRH: H37Rv infected and INH+ RIF treated, InRHD: H37Rv infected and INH+RIF +DFO treated, wpt: weeks post treatment, p values at 95 % confidence interval.

**Supplementary Figure S9:** Pathway enrichment analysis results for treated groups in comparison to infected controls at one and two wpt. wpt: weeks post treatment, p values at 95% confidence interval.

**Supplementary Figure S10: A.** Schematic presentation of the method used for Mtb infection, treatment and intracellular cytokine staining of the lung cells **B.** CD3+ and CD4+ T cell frequency in the infected (In) vs INH+ RIF treated with or without DFO **C.** Percentage of CD4+ T-cells that are IFN-γ, IL-17A and TNF-α positive in the infected (In) vs INH+ RIF treated with or without DFO. n = 5/group/time points for In, InRH group and n = 4 for InRHD group; Results are determined using unpaired, non-parametric two-tailed t-tests. ns: not significant.

**Supplementary Table S1: The list of DEG in DFO-treated Mtb H37Rv compared to control as determined using edgeR in iDEP.96 platform.** The DEGs having FC>2 (log2FC>1), at BH adjusted P-value < 0.05 were considered significant.

**Supplementary Table S2: Expression levels of genes involved in iron metabolism in *Mycobacterium tuberculosis*.** Normalized expression matrix was used for paired, 2-tailed student’s t-test for determination of P-values. The gene set was borrowed from a review by Rodriguez et. al. (Tends in microbiology, 14(7), 2006).

## Notes

### Competing Interest Statement

The authors have declared no competing interest.

### Summary of Updates

We have incorporated the results of transcriptomics and metabolomics of DFO treated Mtb. Also, we have performed an independent experiment to monitor the immune cell distribution in the lungs of Mtb infected C57BL/6 mice receiving either isoniazid and rifampicin or deferoxamine adjunct to isoniazid and rifampicin. We have also replaced the histology images with better resolution ones.

