## Supplemental Information for "Deferoxamine as adjunct therapeutics for tuberculosis"

### Supplementary Figure S1

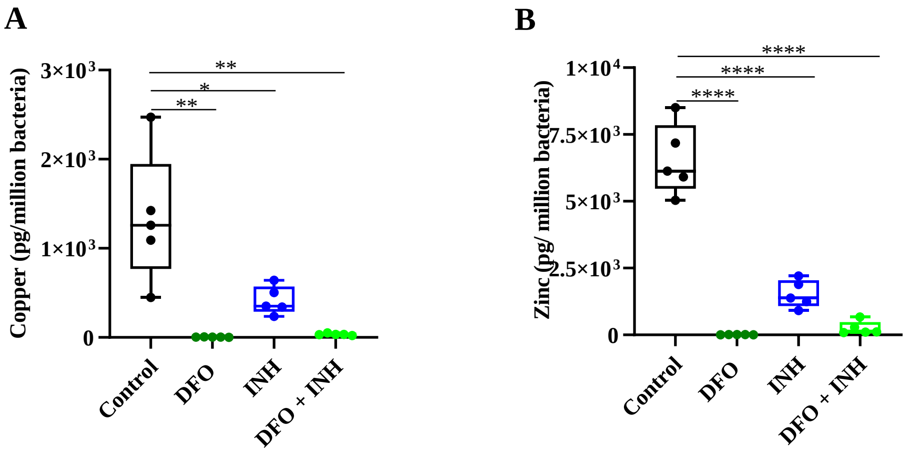


**Supplementary Figure S1: A and B.** Intracellular copper and zinc levels in drug-treated and control Mycobacterial culture at 5th-day post-treatment quantified using ICP-MS. Results are determined using unpaired, non-parametric two-tailed t-tests. n= 5 per group, ns: not significant; *: p<0.05, **: p<0.01; ****: p<0.0001 at 95% confidence interval.

### Supplementary Figure S2

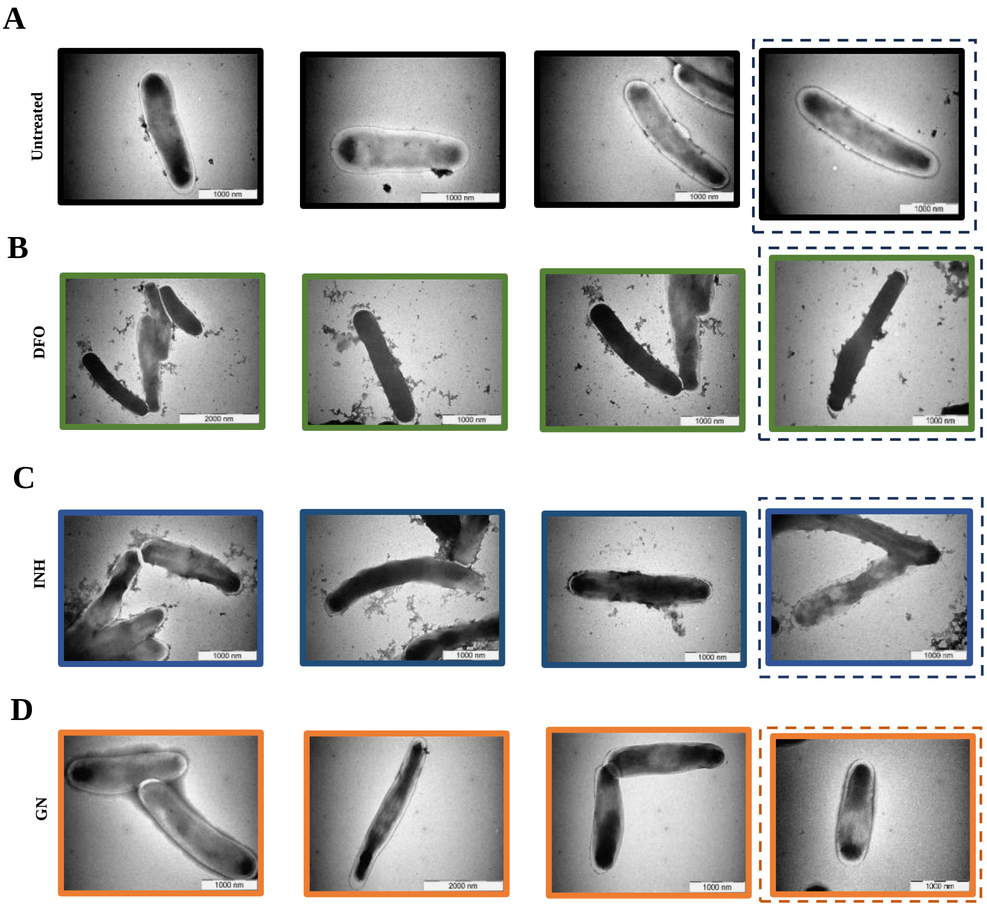


**Supplementary Figure S2:** Transmission electron micrographs of untreated Mtb **(A)**, DFO treated **(B)**, INH treated **(C)**, GN treated **(D)** harvested at 5th day post-drug treatment. Images with dashed boundary are referred in the main figures.

**Supplementary Figure S3**

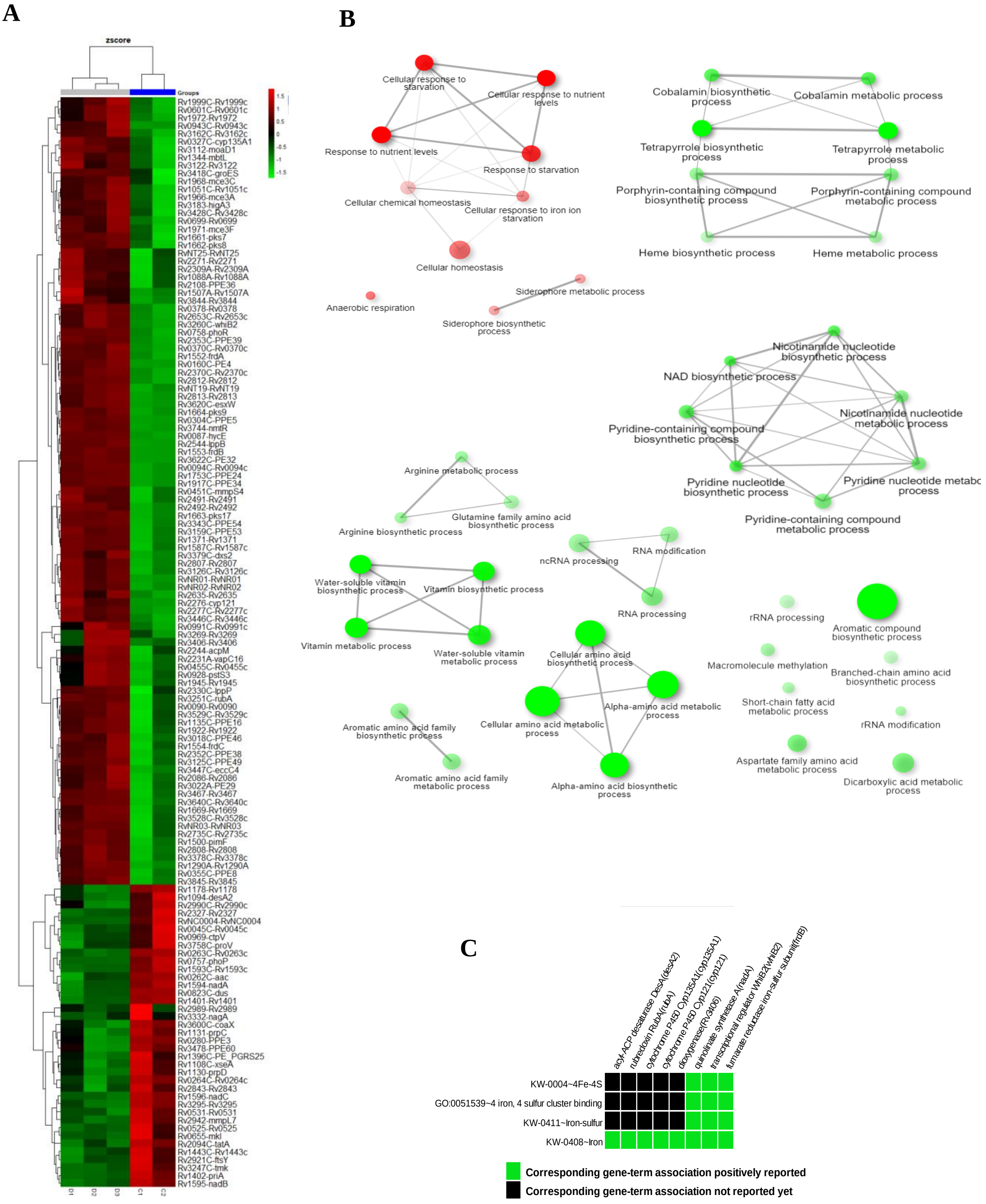


**Supplementary figure S3: DFO alters iron metabolism and various pathways of Mtb H37Rv. A**. Heatmap showing the deregulated (up in red and down in green) genes in the DFO (D1, D2, D3) vs control Mtb (C1, C2) as evident from the Z-score values. **B.** Pathway analysis using Preranked FGSEA in IDEP.96 tool. Red indicates upregulated and green shows downregulated pathways in Mtb. The intensity of the colour and diameter of the node corresponds to the number of genes impacted in a particular pathway. **C.** Iron-sulfur cluster and iron-binding proteins impacted upon DFO treatment. Green shows presence and Black shows absence of the Iron-sulfur/Iron-binding nature of the protein.

**Supplementary Figure S4**

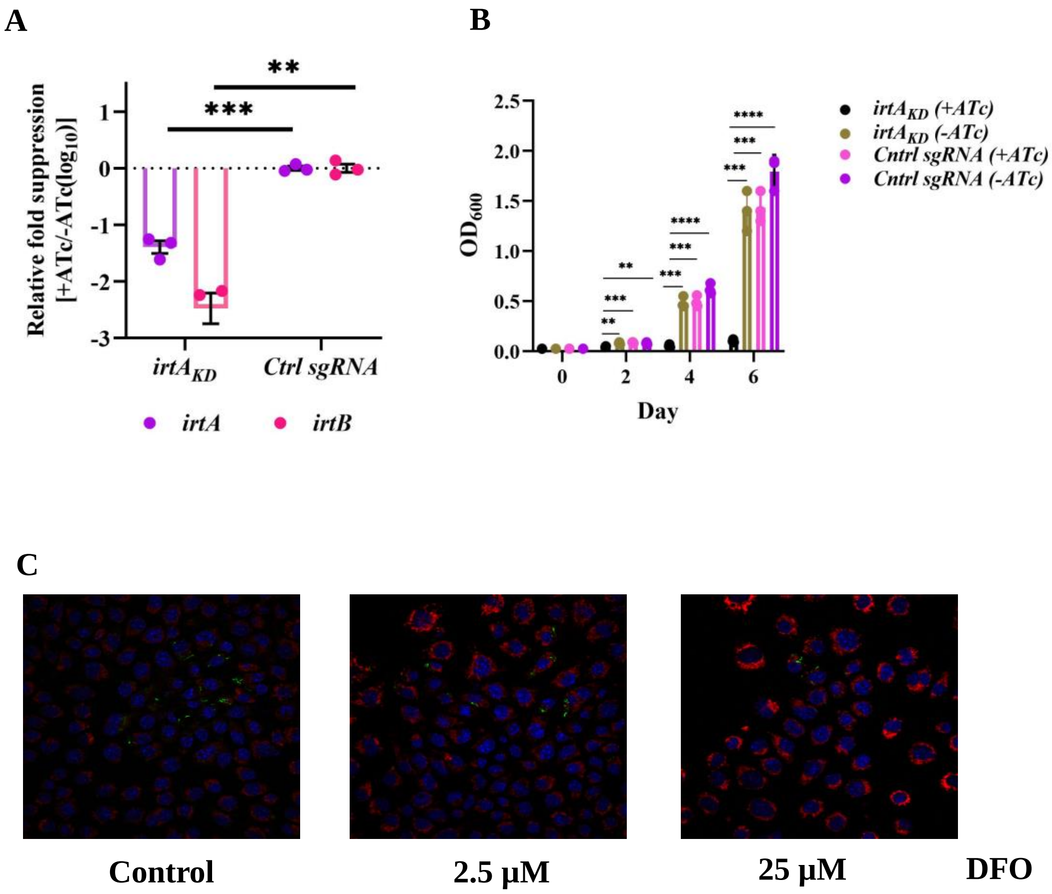


**Supplementary Figure S4: irtA knock down in Mtb inhibits its growth and DFO exposure reduces Mtb growth in J774A1 cells. A.** Relative suppression of *irtA* and *irtB* expression levels in $irtA_{KD}$ and cntrl sgRNA strains (mean ± SEM, $n = 3$ biological replicates). **B.** Relative suppression of *irtA* and *irtB* expression levels in $irtA_{KD}$ and cntrl sgRNA strains **C.** Fluorescent confocal microcopy images acquired at 60X magnification showing the effect of DFO pretreatment on Mtb H37Rv infected J774A.1 murine cells. Blue color represents nuclei stained with DAPI and red color indicates mitochondria stained with MitoTracker red. Results are determined using unpaired, non-parametric two-tailed t-tests **P ≤ 0.005 and ***P ≤ 0.0005, ****P ≤ 0.00005. Data represents mean ± SEM (standard error mean). (KD=knock down, cntrl=control)

### Supplementary Figure S5

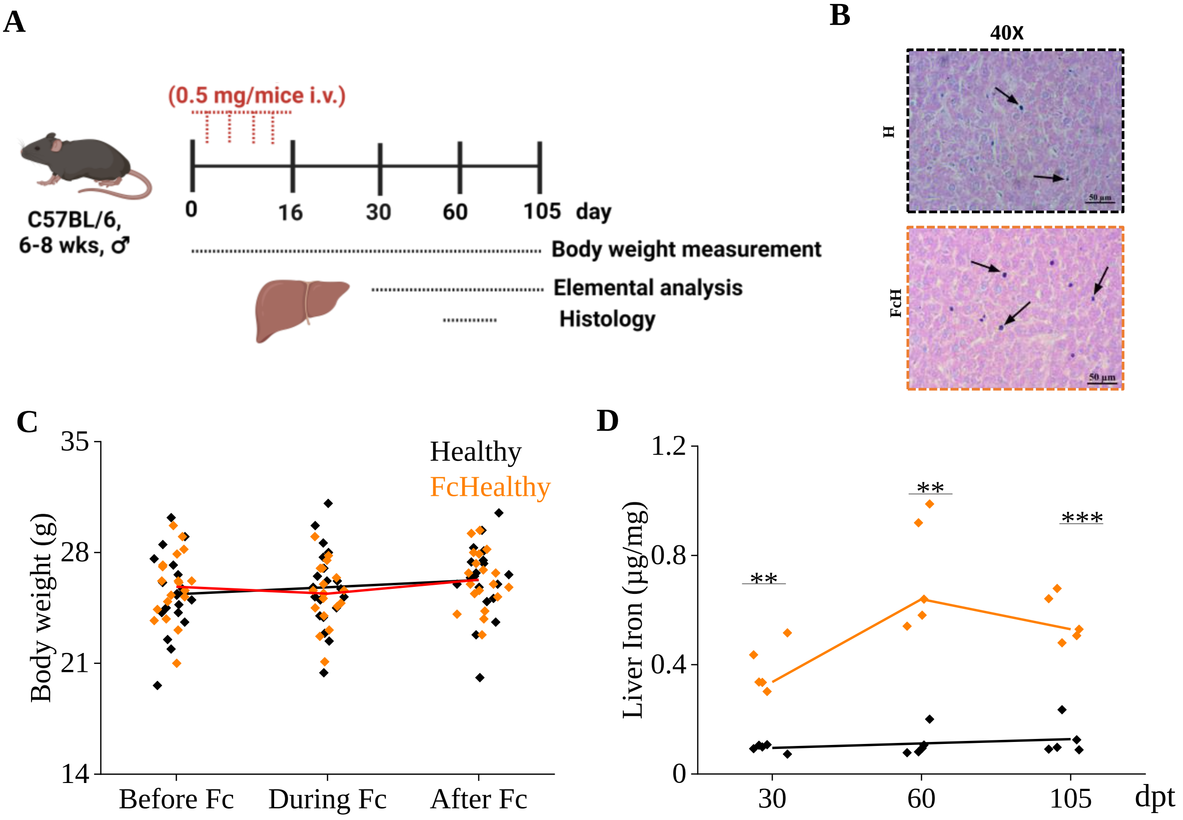


**Supplementary Figure S5: A.** Schematic representation of the method used to create iron loading conditions **B.** Prussian blue staining of the liver tissue at 40X magnification indicating high iron accumulation shown by black arrows in the FcHealthy C57BL/6 mice as compared to healthy controls **C.** Body weight of the C57BL/6 mice before, during and after iron loading for FcHealthy in comparison to healthy controls **D.** Liver iron levels, estimated using ICP-MS at different time points during iron loading as compared to healthy controls. Scale bar for histology images: 50 µm, Fc: ferric carboxymaltose supplemented group. Results are determined using unpaired, non-parametric two-tailed t-tests where ns: not significant; **: $p<0.01$; ***: $p<0.001$ at 95% confidence interval.

### Supplementary Figure S6

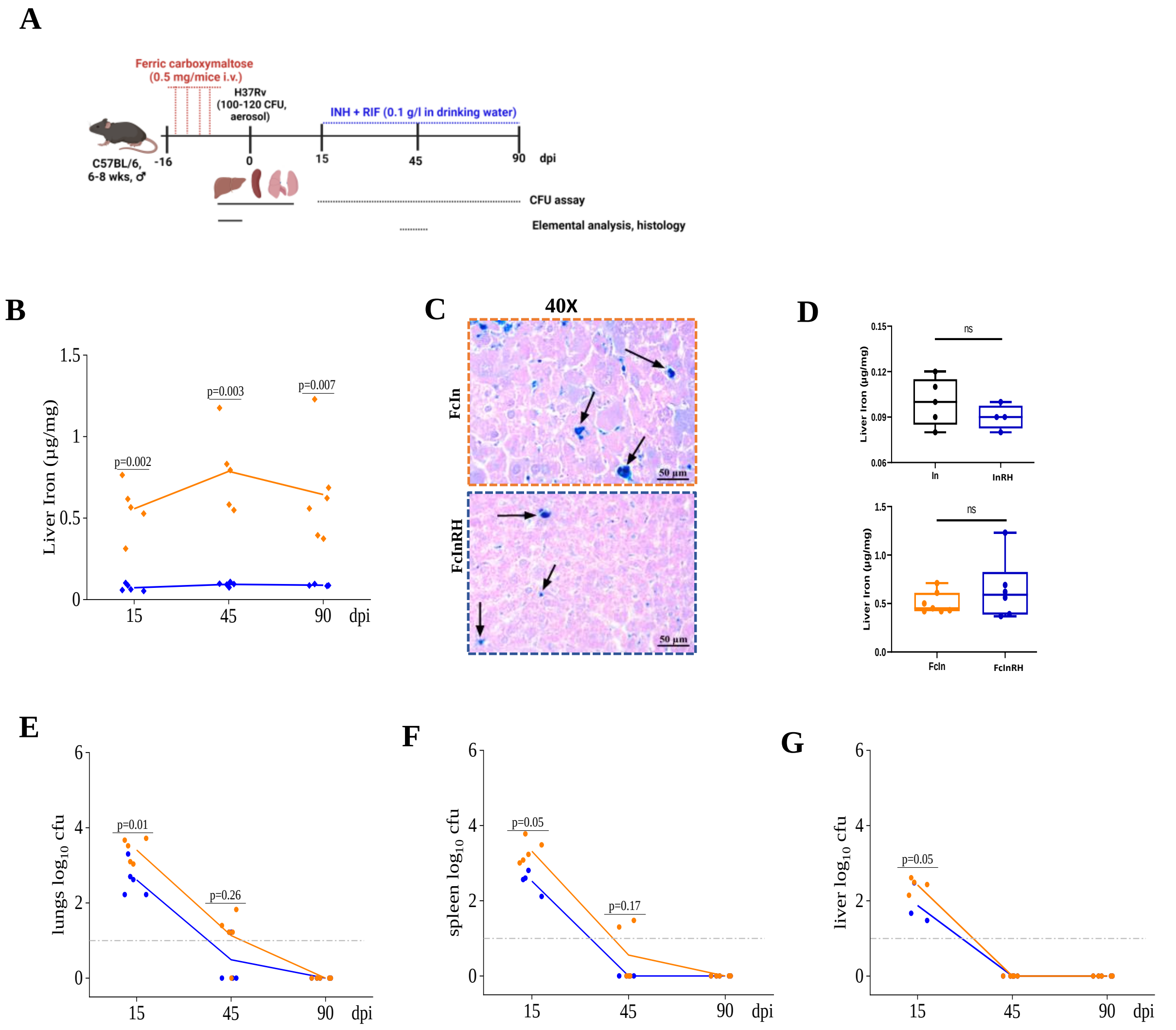


**Supplementary Figure S6: A.** Schematic presentation of the method used to create iron loading condition, Mycobacterial infection and treatment (with isoniazid: INH and rifampicin: RIF) experiment. **B.** Liver iron levels, as estimated using ICP-MS, in iron loaded and control mice groups. **C.** Prussian Staining of liver tissues at 40X magnification indicates high iron accumulation shown by black arrows in iron-loaded mice and did not change significantly upon drug treatment (**D**). Mycobacterial load in the lungs (**E**), spleen (**F**) and liver (**G**) of the *Mycobacteria tuberculosis* H37Rv infected C57BL/6 mice.. Dashed lines in the CFU plots represent the limit of detection. n = 5/group/time points; i.v.: intra-venous injection; dpi: days post-infection; CFU: colony forming units; scale bar for histology images: 50 µm. Results are determined using unpaired, non-parametric two-tailed t-tests where ns: not significant, p values at 95% confidence interval.

**Supplementary Figure S7**

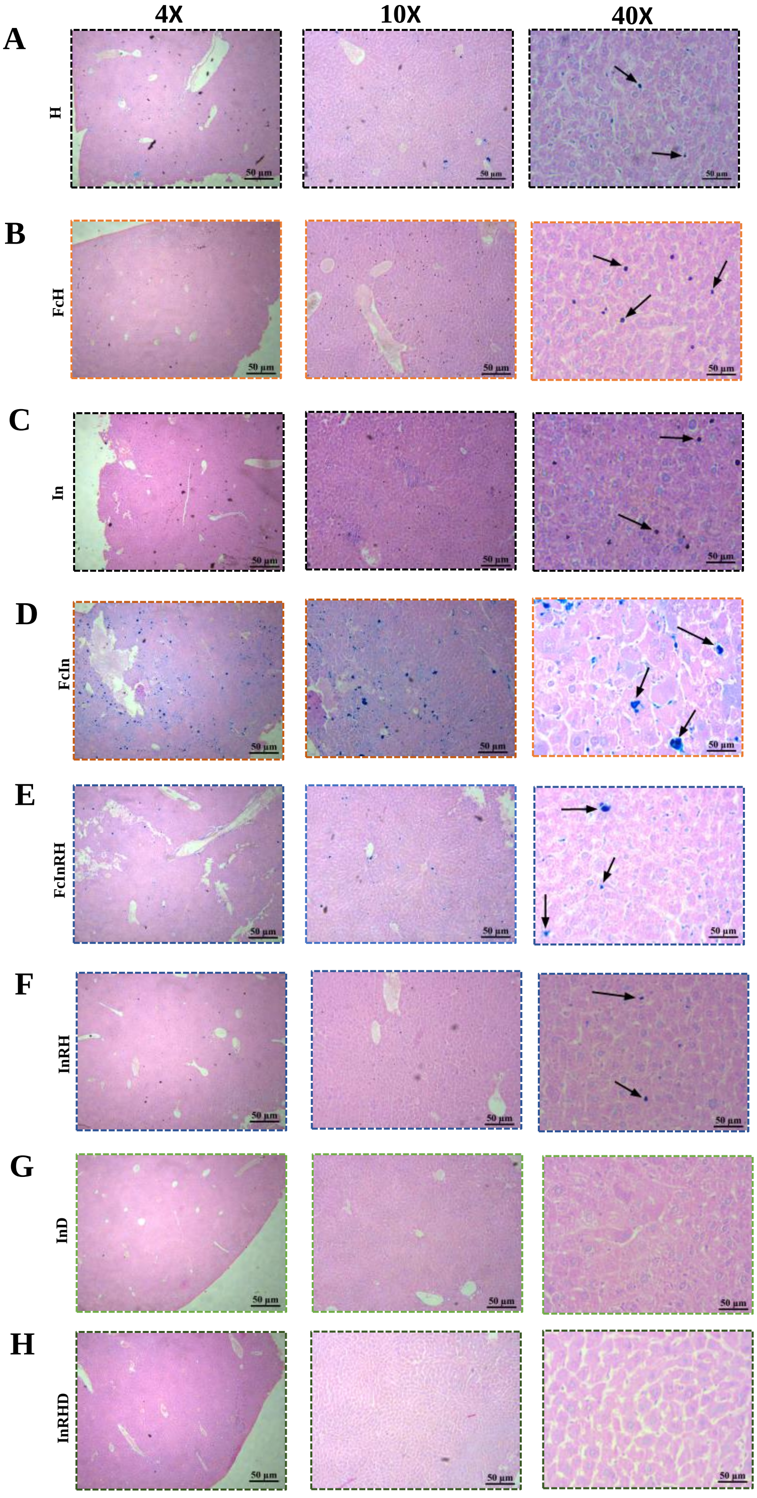


**Supplementary Figure S7:** Histology images of liver tissue stained with Prussian blue staining showing iron accumulation in H (A), FcH (B), In (C), FcIn (D), InRH (E) , FcInRH (F) as compared to minimum iron in InD (G) and InRHD (H) groups at 4X, 10X, 40X magnification respectively. Scale bar: 50µm, In: Mtb infected C57BL/6 mice; FcIn: Mtb infected Fe supplemented C57BL/6 mice; InD: Deferoxamine (DFO) treated Mtb infected mice; InRH: Isoniazid (INH) and Rifampicin (RIF) treated Mtb infected C57BL/6 mice; FcInRH: INH and RIF treated Mtb infected iron supplemented C57BL/6 mice; InRHD: Mtb infected C57BL/6 mice receiving both DFO, INH and RIF; Healthy: Control mice (H); FcHealthy: Control mice with iron supplementation (FcH).

### Supplementary Figure S8

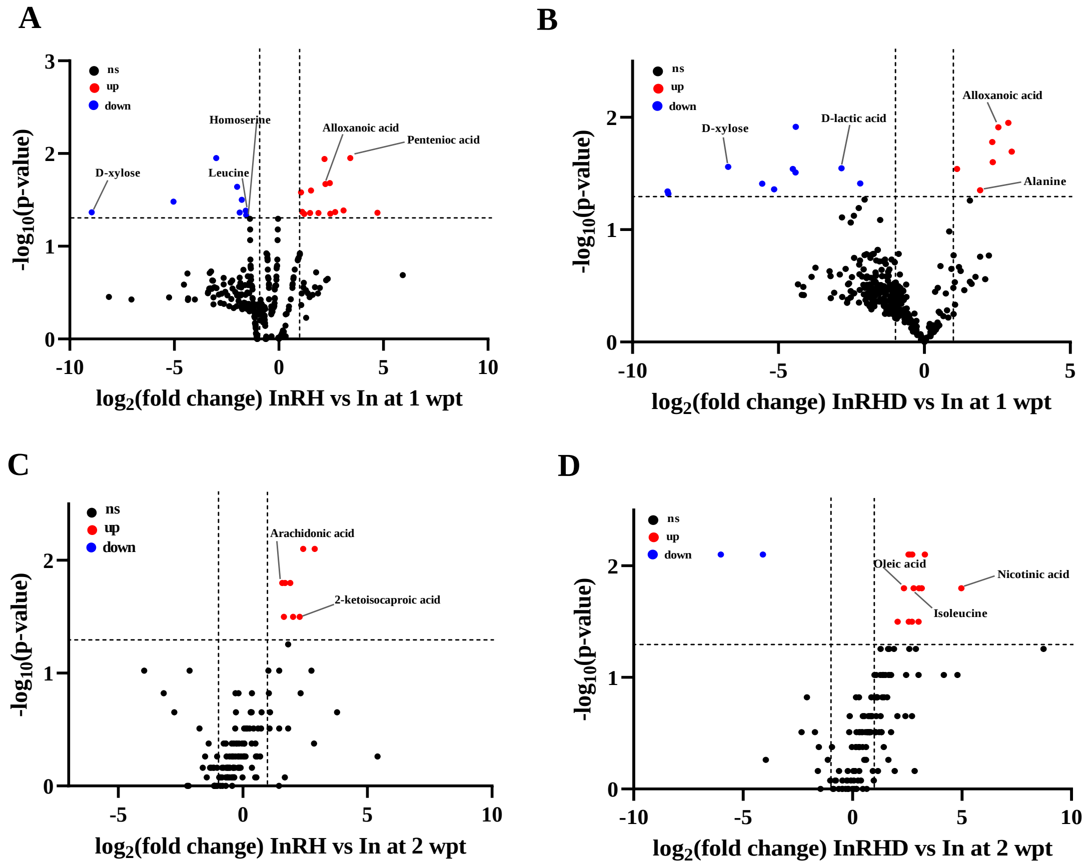


**Supplementary Figure S8:** Liver metabolome changes upon treatment with INH and RIF with or without DFO at 1st and 2nd weeks. InRH vs In **(A)** and InRHD vs In **(B)** InRH vs In **(A)** and InRHD vs In. ns: non significant, up: upregulated, down: down regulated, p value at 95 % confidence interval, In: H37Rv infected control, InRH: H37Rv infected and INH+ RIF treated, InRHD: H37Rv infected and INH+RIF +DFO treated, wpt: weeks post treatment, p values at 95 % confidence interval.

### Supplementary Figure S9

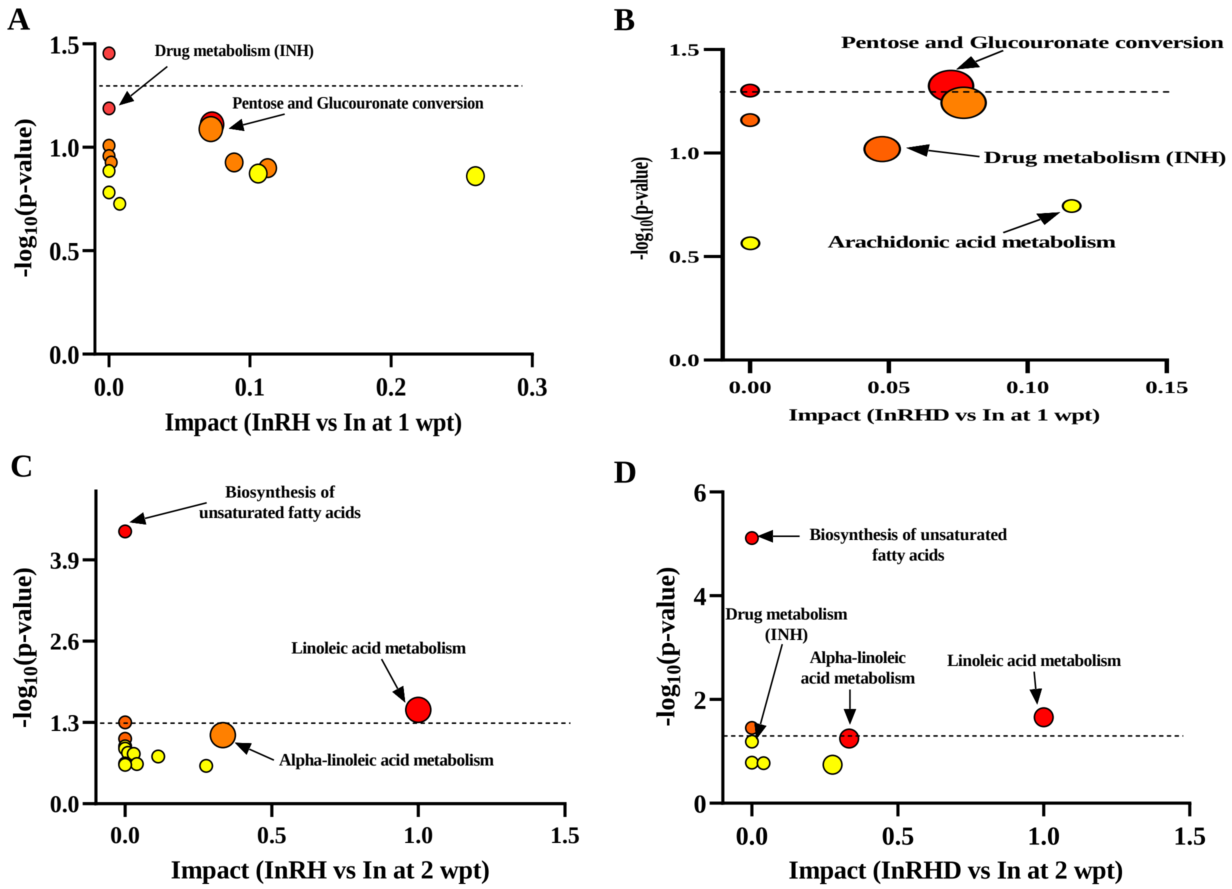


**Supplementary Figure S9:** Pathway enrichment analysis results for treated groups in comparison to infected controls at one and two wpt. wpt: weeks post treatment, p values at 95% confidence interval.

### Supplementary Figure S10

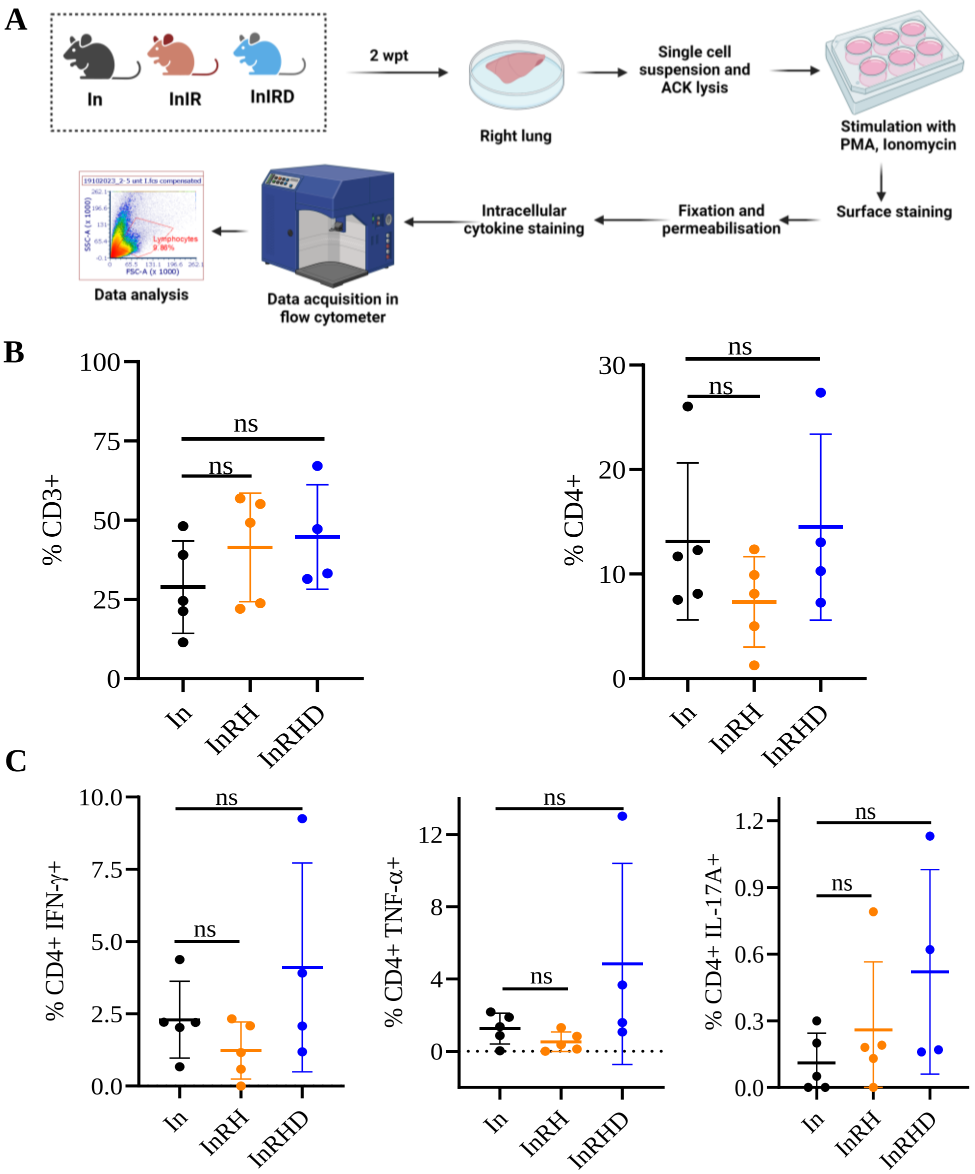


**Supplementary Figure S10: A.** Schematic presentation of the method used for Mtb infection, treatment and intracellular cytokine staining of the lung cells **B.** CD3+ and CD4+ T cell frequency in the infected (In) vs INH+ RIF treated with or without DFO **C.** Percentage of CD4+ T-cells that are IFN-γ, IL-17A and TNF-α positive in the infected (In) vs INH+ RIF treated with or without DFO. n = 5/group/time points for In, InRH group and n=4 for InRHD group; Results are determined using unpaired, non-parametric two-tailed t-tests. ns: not significant.

**Supplementary Table S1: The list of DEG in DFO-treated Mtb H37Rv compared to control as determined using edgeR in iDEP.96 platform.** The DEGs having FC>2 ($\log_2$FC>1), at BH adjusted (Adju.) P-value < 0.05 were considered significant.

| Gene | $\log_2$FC | Adju. P-value |
|---|---|---|
| Rv0758 | 7.24 | <0.0001 |
| Rv1587c | 5.185 | <0.0001 |
| Rv0094c | 4.775 | <0.0001 |
| Rv2353c | 4.731 | <0.0001 |
| Rv3269 | 4.155 | 0.038 |
| Rv2635 | 3.816 | <0.0001 |
| Rv2812 | 3.497 | <0.0001 |
| Rv3406 | 3.482 | 0.027 |
| Rv2813 | 3.246 | <0.0001 |
| Rv3418c | 3.029 | 0.024 |
| Rv1290A | 3.022 | <0.0001 |
| Rvnr03 | 2.999 | <0.0001 |
| Rv1553 | 2.848 | <0.0001 |
| Rvnr01 | 2.843 | <0.0001 |
| Rv0601c | 2.827 | 0.001 |
| Rvnr02 | 2.696 | <0.0001 |
| Rv3467 | 2.667 | <0.0001 |
| Rv1344 | 2.592 | <0.0001 |
| Rv1917c | 2.562 | <0.0001 |
| Rv1371 | 2.54 | <0.0001 |
| Rv3428c | 2.495 | 0.003 |
| Rv1972 | 2.477 | 0.001 |
| Rv3343c | 2.464 | <0.0001 |
| Rv1552 | 2.426 | <0.0001 |
| Rv2544 | 2.415 | <0.0001 |
| Rv2370c | 2.405 | <0.0001 |
| Rv2277c | 2.389 | <0.0001 |
| Rv3528c | 2.386 | 0.001 |
| Rv3162c | 2.376 | <0.001 |
| Rv2244 | 2.376 | 0.046 |
| Rv1554 | 2.361 | 0.003 |
| Rv0699 | 2.354 | 0.001 |
| Rv3022A | 2.283 | 0.002 |
| Rv1663 | 2.252 | <0.0001 |
| Rvnt25 | 2.232 | 0.012 |
| Rv3622c | 2.23 | <0.0001 |
| Rv3112 | 2.142 | 0.001 |
| Rv0304c | 2.115 | <0.0001 |
| Rv2271 | 2.101 | 0.009 |
| Rv1999c | 2.091 | 0.007 |
| Rv3844 | 2.086 | 0.004 |
| Rv3126c | 2.059 | 0.001 |
| Rv1669 | 1.991 | 0.002 |
| Rv1662 | 1.976 | 0.001 |
| Rv2653c | 1.969 | 0.009 |
| Rv2330c | 1.95 | 0.023 |
| Rv2352c | 1.919 | 0.001 |
| Rv3260c | 1.839 | 0.005 |
| Rv1968 | 1.816 | 0.048 |
| Rv2108 | 1.8 | 0.016 |
| Rv0455c | 1.792 | 0.041 |
| Rv3251c | 1.768 | 0.005 |
| Rv1051c | 1.766 | 0.010 |
| Rv0928 | 1.756 | 0.038 |
| Rv0355c | 1.7 | 0.009 |
| Rv1135c | 1.699 | 0.009 |
| Rv3183 | 1.684 | 0.020 |
| Rv0378 | 1.657 | 0.009 |
| Rv1507A | 1.642 | 0.031 |
| Rvnt19 | 1.621 | 0.009 |
| Rv1971 | 1.621 | 0.015 |
| Rv1945 | 1.606 | 0.038 |
| Rv2309A | 1.578 | 0.032 |
| Rv1088a | 1.574 | 0.026 |
| Rv0991c | 1.57 | 0.040 |
| Rv2276 | 1.568 | 0.003 |
| Rv0090 | 1.561 | 0.018 |
| Rv0160c | 1.541 | 0.002 |
| Rv3159c | 1.54 | 0.004 |
| Rv2491 | 1.539 | 0.012 |
| Rv3379c | 1.531 | 0.005 |
| Rv1661 | 1.526 | 0.017 |
| Rv0943c | 1.503 | 0.015 |
| Rv3378c | 1.499 | 0.015 |
| Rv3018c | 1.489 | 0.010 |
| Rv3125c | 1.477 | 0.009 |
| Rv2807 | 1.418 | 0.009 |
| Rv3529c | 1.411 | 0.038 |
| Rv2231A | 1.404 | 0.031 |
| Rv3640c | 1.396 | 0.016 |
| Rv2086 | 1.364 | 0.043 |
| Rv1966 | 1.355 | 0.041 |
| Rv3122 | 1.342 | 0.038 |
| Rv3620c | 1.329 | 0.010 |
| Rv3446c | 1.308 | 0.043 |
| Rv2735c | 1.298 | 0.032 |
| Rv0451c | 1.294 | 0.040 |
| Rv1922 | 1.276 | 0.038 |
| Rv1500 | 1.273 | 0.043 |
| Rv1664 | 1.272 | 0.022 |
| Rv0327c | 1.263 | 0.033 |
| Rv1753c | 1.232 | 0.041 |
| Rv0370c | 1.227 | 0.043 |
| Rv2808 | 1.22 | 0.033 |
| Rv2492 | 1.219 | 0.036 |
| Rv3447c | 1.218 | 0.045 |
| Rv3845 | 1.206 | 0.038 |
| Rv3744 | 1.198 | 0.025 |
| Rv0087 | 1.194 | 0.036 |
| RVnc0004 | -1.201 | 0.031 |
| Rv1443c | -1.226 | 0.045 |
| Rv3247c | -1.263 | 0.025 |
| Rv2921c | -1.266 | 0.041 |
| Rv2327 | -1.269 | 0.028 |
| Rv2094c | -1.286 | 0.027 |
| Rv0531 | -1.297 | 0.029 |
| Rv3758c | -1.313 | 0.043 |
| Rv2843 | -1.345 | 0.049 |
| Rv3295 | -1.369 | 0.007 |
| Rv0525 | -1.382 | 0.024 |
| Rv0823c | -1.425 | 0.005 |
| Rv0655 | -1.43 | 0.034 |
| Rv2942 | -1.542 | 0.009 |
| Rv0969 | -1.599 | 0.015 |
| Rv1108c | -1.606 | 0.015 |
| Rv1402 | -1.635 | 0.002 |
| Rv1178 | -1.666 | 0.002 |
| Rv0045c | -1.668 | 0.012 |
| Rv1401 | -1.682 | <0.001 |
| Rv3600c | -1.815 | 0.002 |
| Rv0262c | -2.006 | <0.0001 |
| Rv0757 | -2.276 | <0.0001 |
| Rv0264c | -2.331 | <0.0001 |
| Rv1594 | -2.391 | <0.0001 |
| Rv0263c | -2.396 | <0.0001 |
| Rv3332 | -2.408 | 0.033 |
| Rv1094 | -2.496 | 0.001 |
| Rv0280 | -2.612 | 0.003 |
| Rv1396c | -2.652 | 0.004 |
| Rv1596 | -2.927 | <0.0001 |
| Rv1595 | -2.958 | <0.0001 |
| Rv1593c | -2.96 | <0.0001 |
| Rv2990c | -3.181 | 0.009 |
| Rv2989 | -3.367 | 0.038 |
| Rv1130 | -3.71 | <0.001 |
| Rv3478 | -3.944 | 0.026 |
| Rv1131 | -4.521 | <0.001 |

**Supplementary Table S2: Expression levels of genes involved in iron metabolism in *Mycobacterium tuberculosis*.** A normalized expression matrix was used for paired, 2-tailed student's t-test for the determination of P-values. The gene set was borrowed from a review by Rodriguez et. al. (Tends in microbiology, 14(7), 2006).

| S. No. | Locus Tag | GeneID | D1 | D2 | D3 | C1 | C2 | $Log_2$(DFO/Control) | P-val |
|---|---|---|---|---|---|---|---|---|---|
| **1** | **Rv3622c** | **PE32** | 3.04 | 3.15 | 3.08 | 0.87 | 0.88 | **1.82** | **0.00001** |
| **2** | **Rv0304c** | **PPE5** | 8.62 | 8.51 | 8.62 | 6.45 | 6.5 | **0.41** | **0.00003** |
| **3** | **Rv1917c** | **PPE34** | 7.81 | 7.95 | 7.95 | 5.25 | 5.43 | **0.57** | **0.0001** |
| **4** | **Rv1753c** | **PPE24** | 7.88 | 7.93 | 7.94 | 6.63 | 6.74 | **0.24** | **0.0001** |
| **5** | **Rv2353c** | **PPE39** | 5.57 | 5.31 | 5.51 | 1.03 | 0.42 | **2.91** | **0.0003** |
| **6** | **Rv0160c** | **PE4** | 5.13 | 5.22 | 5.39 | 3.72 | 3.71 | **0.50** | **0.0006** |
| **7** | **Rv3343c** | **PPE54** | 9.68 | 9.68 | 9.74 | 6.96 | 7.48 | **0.43** | **0.001** |
| **8** | **Rv0354c** | **PPE7** | 6.46 | 6.39 | 6.38 | 5.85 | 5.93 | **0.12** | **0.001** |
| **9** | **Rv3159c** | **PPE53** | 7.11 | 6.88 | 6.94 | 5.23 | 5.62 | **0.36** | **0.003** |
| **10** | **Rv0355c** | **PPE8** | 9.36 | 9.61 | 9.62 | 7.56 | 8.06 | **0.29** | **0.004** |
| **11** | **Rv3125c** | **PPE49** | 5.72 | 5.76 | 5.96 | 4.01 | 4.62 | **0.43** | **0.009** |
| **12** | **Rv3018c** | **PPE46** | 5.22 | 5.09 | 5.5 | 3.51 | 4.03 | **0.48** | **0.009** |
| **13** | **Rv2352c** | **PPE38** | 6.42 | 6.57 | 6.96 | 4.37 | 5.05 | **0.50** | **0.009** |
| **14** | **Rv1348** | **irtA** | 7.2 | 6.95 | 6.88 | 6.32 | 6.2 | **0.16** | **0.011** |
| **15** | **Rv0878c** | **PPE13** | 4.66 | 4.71 | 4.67 | 4.07 | 3.69 | **0.27** | **0.011** |
| **16** | **Rv2408** | **PE24** | 4.06 | 4.02 | 4.23 | 3.51 | 3.64 | **0.20** | **0.012** |
| **17** | **Rv1344** | **mbtL** | 7.15 | 6.38 | 6.7 | 4.62 | 3.56 | **0.72** | **0.012** |
| **18** | **Rv3350c** | **PPE56** | 9.06 | 9.16 | 9.29 | 7.9 | 8.36 | **0.17** | **0.012** |
| **19** | **Rv3022A** | **PE29** | 2.01 | 2.45 | 2.82 | -0.31 | 0.6 | **4.06** | **0.015** |
| **20** | **Rv0151c** | **PE1** | 6.48 | 6.21 | 6.31 | 5.46 | 5.75 | **0.18** | **0.016** |
| **21** | **Rv3245c** | **mtrB** | 7.82 | 7.94 | 7.94 | 8.25 | 8.46 | **-0.08** | **0.016** |
| **22** | **Rv0256c** | **PPE2** | 7.6 | 7.75 | 7.76 | 8.18 | 8.49 | **-0.11** | **0.017** |
| **23** | **Rv1135c** | **PPE16** | 5.72 | 6 | 6.11 | 3.68 | 4.66 | **0.51** | **0.02** |
| **24** | **Rv2123** | **PPE37** | 4.33 | 4.63 | 4.68 | 3.97 | 3.91 | **0.21** | **0.023** |
| **25** | **Rv0754** | **PE_PGRS11** | 5.18 | 5.19 | 5.57 | 4.66 | 4.52 | **0.21** | **0.025** |
| **26** | **Rv2108** | **PPE36** | 7.98 | 7.29 | 7.19 | 5.24 | 6.1 | **0.40** | **0.027** |
| **27** | **Rv3812** | **PE_PGRS62** | 6.57 | 6.9 | 6.87 | 6.09 | 6.27 | **0.13** | **0.028** |
| **28** | **Rv0280** | **PPE3** | 6.13 | 4.3 | 4.47 | 7.99 | 7.65 | **-0.65** | **0.033** |
| **29** | **Rv0755c** | **PPE12** | 7.84 | 8.55 | 8.52 | 6.46 | 7.26 | **0.28** | **0.041** |
| **30** | **Rv3595c** | **PE_PGRS59** | 3.96 | 4.73 | 4.7 | 5.57 | 5.48 | **-0.31** | **0.047** |
| 31 | Rv1346 | mbtN | 7.59 | 7.39 | 7.36 | 6.86 | 6.05 | 0.21 | 0.05 |
| 32 | Rv0159c | PE3 | 5.89 | 5.91 | 6.07 | 5.75 | 5.69 | 0.06 | 0.05 |
| 33 | Rv1396c | PE_PGRS25 | 3.63 | 2.33 | 2.6 | 6.26 | 4.41 | -0.90 | 0.06 |
| 34 | Rv0578c | PE_PGRS7 | 5.9 | 6.6 | 6.62 | 7.28 | 7.22 | -0.19 | 0.06 |
| 35 | Rv1349 | irtB | 6.38 | 6.36 | 6.52 | 6.15 | 5.71 | 0.11 | 0.07 |
| 36 | Rv0978c | PE_PGRS17 | 2.52 | 2.69 | 2.47 | 2.35 | 2.17 | 0.18 | 0.07 |
| 37 | Rv0152c | PE2 | 6.38 | 5.71 | 5.87 | 5.16 | 5.35 | 0.19 | 0.07 |
| 38 | Rv1196 | PPE18 | 8.67 | 7.34 | 7.42 | 9.63 | 9.15 | -0.27 | 0.07 |
| 39 | Rv1361c | PPE19 | 8.03 | 6.52 | 6.68 | 8.98 | 8.23 | -0.28 | 0.10 |
| 40 | Rv1548c | PPE21 | 6.87 | 6.94 | 6.99 | 4.75 | 6.4 | 0.31 | 0.11 |
| 41 | Rv0305c | PPE6 | 7.7 | 8.44 | 8.44 | 6.9 | 7.65 | 0.17 | 0.11 |
| 42 | Rv0742 | PE_PGRS8 | 2.29 | 1.58 | 1.61 | 3.5 | 2.3 | -0.67 | 0.14 |
| 43 | Rv1791 | PE19 | 8.98 | 9.42 | 9.35 | 7.56 | 8.87 | 0.17 | 0.14 |
| 44 | Rv1789 | PPE26 | 6.76 | 7.03 | 7.06 | 7.34 | 7.13 | -0.06 | 0.14 |
| 45 | Rv0977 | PE_PGRS16 | 6.14 | 6.12 | 6.18 | 5.65 | 6.07 | 0.07 | 0.16 |
| 46 | Rv1347c | mbtK | 5.94 | 5.2 | 5.25 | 5.1 | 4.05 | 0.26 | 0.17 |
| 47 | Rv1195 | PE13 | 8.97 | 7.34 | 7.37 | 9.18 | 9 | -0.20 | 0.18 |
| 48 | Rv1800 | PPE28 | 6.26 | 6.83 | 6.87 | 5.75 | 6.4 | 0.13 | 0.19 |
| 49 | Rv0278c | PE_PGRS3 | 2.62 | 3.14 | 3.3 | 3.81 | 3.3 | -0.24 | 0.19 |
| 50 | Rv1387 | PPE20 | 9.02 | 6.02 | 5.93 | 9.08 | 9.13 | -0.38 | 0.20 |
| 51 | Rv1325c | PE_PGRS24 | 3.62 | 3.61 | 3.71 | 4.43 | 3.72 | -0.16 | 0.20 |
| 52 | Rv1168c | PPE17 | 6.76 | 8.03 | 7.96 | 6.58 | 6.93 | 0.17 | 0.22 |
| 53 | Rv1088 | PE9 | 2.1 | 1.91 | 2.3 | 2.01 | 1.02 | 0.47 | 0.23 |
| 54 | Rv1305 | atpE | 7.39 | 7.47 | 7.28 | 6.31 | 7.33 | 0.11 | 0.24 |
| 55 | Rv1305 | atpE | 7.39 | 7.47 | 7.28 | 6.31 | 7.33 | 0.11 | 0.24 |

| | | | | | | | | | |
|---|---|---|---|---|---|---|---|---|---|
| **56** | Rv1345 | mbtM | 6.45 | 6.81 | 6.91 | 6.66 | 5.46 | 0.15 | 0.26 |
| **57** | Rv0286 | PPE4 | 8.19 | 7.65 | 7.63 | 8.81 | 7.94 | -0.10 | 0.26 |
| **58** | Rv1039c | PPE15 | 3.91 | 4.38 | 4.69 | 4.45 | 5.73 | -0.23 | 0.26 |
| **59** | Rv0096 | PPE1 | 4.33 | 4.62 | 5.06 | 4.58 | 3.09 | 0.28 | 0.27 |
| **60** | Rv0109 | PE_PGRS1 | 3.47 | 4.27 | 4.43 | 3.23 | 3.83 | 0.20 | 0.32 |
| **61** | Rv1040c | PE8 | 1.74 | 3.27 | 3.19 | 2.82 | 5.31 | -0.57 | 0.32 |
| **62** | Rv1803c | PE_PGRS32 | 5.24 | 6.11 | 6.06 | 4.93 | 5.64 | 0.13 | 0.33 |
| **63** | Rv0916c | PE7 | 3.38 | 4.45 | 4.43 | 3.04 | 3.87 | 0.24 | 0.33 |
| **64** | Rv0746 | PE_PGRS9 | 0.05 | 2.1 | 2.1 | 2.91 | 2.05 | -0.81 | 0.33 |
| **65** | Rv1172c | PE12 | 6.94 | 6.78 | 6.82 | 5.96 | 6.92 | 0.09 | 0.34 |
| **66** | Rv1651c | PE_PGRS30 | 6.07 | 6.17 | 6.37 | 6.66 | 6.2 | -0.05 | 0.35 |
| **67** | Rv0532 | PE_PGRS6 | 2.88 | 3.47 | 3.57 | 3.26 | 2.45 | 0.21 | 0.35 |
| **68** | Rv0980c | PE_PGRS18 | 7.83 | 6.32 | 6.19 | 7.77 | 7.3 | -0.15 | 0.36 |
| **69** | Rv0124 | PE_PGRS2 | 3.84 | 3.12 | 3.43 | 3.86 | 3.66 | -0.12 | 0.36 |
| **70** | Rv1801 | PPE29 | 3.58 | 4.98 | 5.4 | 4.1 | 3.74 | 0.25 | 0.38 |
| **71** | Rv1441c | PE_PGRS26 | 2.83 | 3.5 | 3.77 | 3.53 | 4.01 | -0.16 | 0.39 |
| **72** | Rv0833 | PE_PGRS13 | 4.91 | 5.56 | 5.53 | 6.87 | 5.07 | -0.16 | 0.44 |
| **73** | Rv1386 | PE15 | 7.36 | 5.55 | 5.9 | 8.34 | 6.12 | -0.21 | 0.44 |
| **74** | Rv0832 | PE_PGRS12 | 2.37 | 3.3 | 3.27 | 4.38 | 2.78 | -0.26 | 0.46 |
| **75** | Rv1802 | PPE30 | 3.98 | 4.09 | 4.49 | 4.41 | 4.28 | -0.05 | 0.49 |
| **76** | Rv1705c | PPE22 | 4.11 | 5.01 | 5.18 | 4.27 | 6.85 | -0.22 | 0.50 |
| **77** | Rv1646 | PE17 | 7.19 | 8.3 | 8.21 | 8.11 | 8.39 | -0.06 | 0.51 |
| **78** | Rv1307 | atpH | 9.11 | 8.3 | 8.21 | 9.15 | 8.56 | -0.05 | 0.51 |
| **79** | Rv1430 | PE16 | 6.44 | 7.04 | 7.06 | 6.02 | 7.03 | 0.07 | 0.53 |
| **80** | Rv1806 | PE20 | 3.87 | 7.96 | 7.85 | 2.61 | 7.15 | 0.43 | 0.53 |
| **81** | Rv0915c | PPE14 | 2.35 | 3.18 | 3.6 | 2.36 | 3.03 | 0.18 | 0.56 |
| **82** | Rv0872c | PE_PGRS15 | 4.14 | 6.09 | 6.25 | 6.62 | 5.58 | -0.15 | 0.57 |
| **83** | Rv1068c | PE_PGRS20 | 2.92 | 2.3 | 2.65 | 2.82 | 1.91 | 0.15 | 0.57 |
| **84** | Rv0285 | PE5 | 2.31 | 2.68 | 3.19 | 5.23 | 1.85 | -0.38 | 0.57 |
| **85** | Rv1309 | atpG | 8.08 | 7.49 | 7.38 | 7.71 | 7.94 | -0.03 | 0.59 |
| **86** | Rv1067c | PE_PGRS19 | 0.49 | 0.05 | 1.08 | 1.51 | 0.26 | -0.71 | 0.60 |
| **87** | Rv1243c | PE_PGRS23 | 1.41 | 1.1 | 2.29 | 1.99 | 0.36 | 0.45 | 0.61 |
| **88** | Rv1787 | PPE25 | 5.29 | 6.19 | 6.22 | 5.42 | 5.91 | 0.06 | 0.62 |
| **89** | Rv1768 | PE_PGRS31 | 4.79 | 6.14 | 6.15 | 5.7 | 5.07 | 0.08 | 0.65 |
| **90** | Rv1311 | atpC | 7.86 | 6.86 | 6.72 | 8.17 | 6.77 | -0.06 | 0.67 |
| **91** | Rv1306 | atpF | 8.77 | 7.95 | 7.69 | 8.32 | 8.29 | -0.03 | 0.71 |
| **92** | Rv1310 | atpD | 8.53 | 7.82 | 7.73 | 8.66 | 7.76 | -0.03 | 0.72 |
| **93** | Rv1706c | PPE23 | 4.08 | 5.94 | 5.96 | 3.91 | 7.89 | -0.15 | 0.75 |
| **94** | Rv1452c | PE_PGRS28 | 0.78 | 1.26 | 2.05 | 2.13 | 0.08 | 0.30 | 0.79 |
| **95** | Rv1468c | PE_PGRS29 | 3.32 | 4.7 | 4.91 | 4.84 | 3.35 | 0.07 | 0.81 |
| **96** | Rv0335c | PE6 | 2.26 | 3.37 | 3.39 | 4.03 | 2.37 | -0.09 | 0.82 |
| **97** | Rv1809 | PPE33 | 6.67 | 9.39 | 9.31 | 6.84 | 9.36 | 0.06 | 0.82 |
| **98** | Rv0453 | PPE11 | 4.7 | 5.24 | 5.43 | 3.75 | 7.1 | -0.08 | 0.82 |
| **99** | Rv0834c | PE_PGRS14 | 6.15 | 7.16 | 7.14 | 7.25 | 6.6 | -0.02 | 0.84 |
| **100** | Rv1788 | PE18 | 3.32 | 3.95 | 4.09 | 4.4 | 3.38 | -0.04 | 0.84 |
| **101** | Rv1214c | PE14 | 4.33 | 3.89 | 4.13 | 4.61 | 3.76 | -0.02 | 0.86 |
| **102** | Rv1091 | PE_PGRS22 | 2.42 | 3.2 | 3.4 | 3.76 | 2.48 | -0.05 | 0.86 |
| **103** | Rv0279c | PE_PGRS4 | 3.1 | 3.71 | 3.65 | 3.35 | 3.54 | 0.02 | 0.88 |
| **104** | Rv1808 | PPE32 | 5.81 | 8.69 | 8.61 | 6.66 | 8.34 | 0.04 | 0.89 |
| **105** | Rv1450c | PE_PGRS27 | 1.97 | 2.62 | 2.65 | 3.2 | 1.8 | -0.05 | 0.89 |
| **106** | Rv0442c | PPE10 | 6.16 | 6.28 | 6.29 | 5.95 | 6.6 | -0.01 | 0.90 |
| **107** | Rv1790 | PPE27 | 4.07 | 5.96 | 6.04 | 4.56 | 5.94 | 0.03 | 0.91 |
| **108** | Rv1087 | PE_PGRS21 | 3.64 | 4.63 | 4.61 | 4.84 | 3.86 | -0.02 | 0.92 |
| **109** | Rv0297 | PE_PGRS5 | 4.99 | 6.49 | 6.67 | 6.31 | 5.91 | -0.01 | 0.93 |
| **110** | Rv1308 | atpA | 10.24 | 9.62 | 9.46 | 10.06 | 9.47 | 0.00 | 0.98 |
| **111** | Rv1304 | atpB | 7 | 6.69 | 6.61 | 6.4 | 7.14 | 0.00 | 0.99 |
| **112** | Rv0747 | PE_PGRS10 | 2.81 | 5.46 | 5.49 | 3.23 | 5.95 | 0.00 | 0.99 |
